# Genome-wide screen overexpressing mycobacteriophage Amelie genes identifies multiple inhibitors of mycobacterial growth

**DOI:** 10.1101/2024.08.14.607960

**Authors:** Chelsea Tafoya, Brandon Ching, Elva Garcia, Alyssa Lee, Melissa Acevedo, Kelsey Bass, Elizabeth Chau, Heidi Lin, Kaitlyn Mamora, Michael Reeves, Madyllyne Vaca, William van Iderstein, Luis Velasco, Vivianna Williams, Grant Yonemoto, Tyler Yonemoto, Danielle M. Heller, Arturo Diaz

## Abstract

The genome sequences of thousands of bacteriophages have been determined and functions for many of the encoded genes have been assigned based on homology to characterized sequences. However, functions have not been assigned to more than two-thirds of the identified phage genes as they have no recognizable sequence features. Recent genome-wide overexpression screens have begun to identify bacteriophage genes that encode proteins that reduce or inhibit bacterial growth. This study describes the construction of a plasmid-based overexpression library of 76 genes encoded by Cluster K1 mycobacteriophage Amelie, which is genetically similar to Cluster K phages Waterfoul and Hammy recently described in similar screens and closely related to phages that infect clinically important mycobacteria. 26 out of the 76 genes evaluated in our screen, encompassing 34% of the genome, reduced growth of the host bacterium *Mycobacterium smegmatis* to various degrees. More than one-third of these 26 toxic genes have no known function, and 10 of the 26 genes almost completely abolished host growth upon overexpression. Notably, while several of the toxic genes identified in Amelie shared homologs with other Cluster K phages recently screened, this study uncovered eight previously unknown gene families that exhibit cytotoxic properties, thereby broadening the repertoire of known phage-encoded growth inhibitors. This work, carried out under the HHMI-supported SEA-GENES project (Science Education Alliance Gene-function Exploration by a Network of Emerging Scientists), underscores the importance of comprehensive overexpression screens in elucidating genome-wide patterns of phage gene function and novel interactions between phages and their hosts.

## Introduction

The bacteriophage population is incredibly diverse, with an estimated 10^31^ particles in the biosphere (Hatfull and Hendrix 2011; Hatfull 2015). One way to identify viral diversity is through metagenomics of total concentrated phages collected from environmental samples. This approach generates vast amounts of DNA fragments sequenced at random, but it is difficult to obtain biological materials for further experimentation (Edwards and Rohwer 2005; Kristensen *et al*. 2010). A second approach involves analyzing individually isolated phages using genetic, biochemical and microbiological techniques. The genome sequences of nearly ∼5,000 phages isolated on actinobacterial species have been determined and are characterized by a mosaic composition as a result of horizontal gene transfer (Pedulla *et al*. 2003; Russell and Hatfull 2017). Comparative analyses of the sequenced genomes have revealed more than 30,000 phamilies, or groupings of gene products sharing >25% amino acid identity, of which ∼75% cannot be assigned a function based on homology to characterized sequences (Cresawn *et al*. 2011; Pope *et al*. 2015; Russell and Hatfull 2017; Hatfull 2020; Gauthier *et al*. 2022). In addition to the genes that encode for structural proteins as well those involved in DNA replication, phage genomes harbor a variable array of non-core genes, the vast majority of which have no known function (NKF) (Hatfull 2015). Elucidating the functions of these genes could shed light on the ways that phages manipulate their bacterial hosts as well as help identify bacterial defense mechanisms (Ko and Hatfull 2018, 2020; Hampton *et al*. 2020; Srikant *et al*. 2022). Understanding the interplay between phages and their bacterial hosts has implications for phage-based therapeutic interventions, microbial evolution, and the potential development of new biotechnology tools and applications.

By connecting genotype to phenotype, screens of phage gene overexpression libraries serve as a crucial first step in understanding the role of uncharacterized phage genes. For example, inhibition of bacterial growth upon phage gene overexpression can help identify proteins that interact with host factors (Rybniker *et al*. 2008, 2011; Ko and Hatfull 2018, 2020). Recent overexpression screens done as part of the Science Education Alliance Gene-function Exploration by a Network of Emerging Scientists (SEA-GENES project)(Heller and Sivanathan 2022; Heller *et al*. 2024) demonstrated that anywhere from 25% to one-third of the genes encoded in a single mycobacteriophage genome inhibited *Mycobacterium smegmatis* growth to various degrees (Heller *et al*. 2022; Amaya *et al*. 2023; Pollenz *et al*. 2024). Similarly, a previous study in which close to 200 unrelated genes from 13 diverse phages were tested for cytotoxicity in *M. smegmatis* showed that 23% inhibited growth when overexpressed (Ko and Hatfull 2020). Given the size and diversity of the phage population, further phenotypic exploration of sequenced phage genomes will lead to the identification of more genes that disrupt or inhibit bacterial growth.

In this study we present findings from a genome-wide overexpression screen focusing on phage Amelie, further expanding on the screens conducted through the SEA-GENES program.

Amelie is a temperate siphovirus isolated on *M. smegmatis* that encodes for 77 predicted protein coding genes, of which only 42 can be assigned a predicted function. Through comparative genomic analysis, Amelie is classified within Cluster K (Subcluster K1) and shares genetic similarities with Waterfoul (Subcluster K5) and Hammy (Subcluster K6), for which overexpression screens were recently reported (Heller *et al*. 2022; Amaya *et al*. 2023). Gene content similarity (GCS), which is the number of gene phamilies (phams) shared between the two genomes, divided by the total number of phams in the genome, shows that Amelie is 49.6% similar to Waterfoul and 67.3% similar to Hammy. Additionally, Amelie is related to ZoeJ (Subcluster K2), a phage that was used to treat a patient with drug resistant *Mycobacerium abscessus* (Dedrick *et al*. 2019a, 2019b) and Adephagia (Subcluster K1), which has been shown to infect strains of *Mycobacterium tuberculosis* (Jacobs-Sera *et al*. 2012; Guerrero-Bustamante *et al*. 2021). 26 out of the 76 genes evaluated in our screen inhibited growth to various degrees, encompassing 34% of the genome. Nine of the cytotoxic genes encoded by Amelie are related to genes previously identified as toxic in both Waterfoul and Hammy, seven genes that conferred toxicity are shared by Amelie and Hammy, two toxic genes are shared by only Amelie and Waterfoul, and eight genes are distinct from those identified in Waterfoul and Hammy.

These results highlight the shared and unique genes found within closely related phages to modulate their bacterial hosts.

## Materials and Methods

### Growth of mycobacteria and mycobacteriophage

*Mycobacterium smegmatis mc*^*2*^*155* was grown at 37°C in Middlebrook 7H9 (Difco) broth supplemented with 10% AD (2% w/v Dextrose, 145 mM NaCl, 5% w/v Albumin Fraction V), 0.05% Tween80, and 10 µg/ml cycloheximide (CHX) or on Middlebrook 7H10 (Difco) agar supplemented with 10% AD and 10 µg/ml CHX. 20-200 ng of pExtra plasmid was used to transform M. smegmatis mc^2^155, electrocompetent cells were electroporated, recovered in 7H9 broth for 1.5 h at 37°C with shaking, and transformants selected on 7H10 agar supplemented with 10 µg/ml Kanamycin (GoldBio). Plates were incubated at 37°C for 5 days, at which point colonies were harvested for the cytotoxicity assay. Amelie was propagated on M. smegmatis mc^2^155 grown at 37°C in the presence of 1 mM CaCl_2_ and no Tween in Middlebrook media and top agar.

### Construction of the pExTra Hammy Library

Each Amelie gene was cloned into the pExTra shuttle vector (Heller *et al*. 2022) downstream of an anhydrotetracycline-inducible promoter, *pTet* (Ehrt 2005), to control gene expression, and upstream of linked *mcherry* transcriptional reporter (Fig. 1a). Individual Amelie genes were PCR-amplified using gene-specific primers (Integrated DNA Technologies), Q5 DNA polymerase (New England Biolabs Q5 HotStart 2X Master Mix), and a high-titer Amelie lysate. Forward primers are complementary to the first 15-25 bp of each gene sequence and introduce a uniform ATG start codon. Reverse primers are complementary to the last 15-25 bp of each gene sequence and introduce a uniform TGA codon (Supplementary Table 1). Each primer also contained sequence of homology to the pExtra01 plasmid; forward primers contain a uniform ribosome binding site (RBS) and 5′ 21 bp of homology to pExTra01 downstream of the *pTet* promoter, and all reverse primers contain a separate 5′ 25 bp of homology to pExTra01 upstream of *mcherry* (Supplementary Table 1). Divergent primers pExTra_F and pExTra_R were used to generate linearized pExTra plasmid via PCR (NEB Q5 HotStart 2X Master Mix) of pExTra01 (Heller *et al*. 2022) and assembled with each gene insert by isothermal assembly (NEB HiFi 2X Master Mix). Despite several attempts, pExtra clones containing gene *75* could not be recovered. Recombinant plasmids were recovered by transformation of *E. coli NEB5α F’I*^*Q*^ (New England Biolabs) and selection on LB agar supplemented with 50 µg/ml Kanamycin. The inserted genes for all recovered pExTra plasmids were sequence-verified by Sanger sequencing (Genewiz/Azenta) using sequencing primers pExTra_universalR and pExTra_seqF; longer genes were also sequenced with internal sequencing primers listed in Supplemental Table 1. All plasmid inserts were found to match the published genome sequence.

**Table 1.** Amelie genes observed to inhibit *Mycobacterium smegmatis* growth upon overexpression.

| Amelie gene | Phamily<sup>a</sup> | Length (aa) | Predicted Function | Cytotoxicity<sup>b</sup> |
| --- | --- | --- | --- | --- |
| 8 | 166542 | 182 | Scaffolding protein | 1 |
| 16 | 166435 | 274 | Tail assembly chaperone | 1 |
| 46 | 166376 | 89 | WhiB family transcription factor | 1 |
| 55 | 166327 | 81 | NrdH-like glutaredoxin | 1 |
| 56 | 166434 | 123 | HNH endonuclease | 1 |
| 58 | 164894 | 249 | RusA-like resolvase | 1 |
| 63 | 6240 | 166 | NKF<sup>c</sup> | 1 |
| 65 | 84966 | 289 | NKF; TMD<sup>d</sup> | 1 |
| 74 | 87639 | 139 | NKF | 1 |
| 15 | 166425 | 138 | Tail assembly chaperone | 2 |
| 20 | 166421 | 158 | Minor tail protein | 2 |
| 31 | 166375 | 378 | MRE11 double-stranded break endo/exonuclease | 2 |
| 32 | 95039 | 137 | NKF; HTH DNA binding protein | 2 |
| 50 | 166417 | 295 | Cas4-like exonuclease | 2 |
| 62 | 158927 | 90 | NKF | 2 |
| 9 | 164750 | 306 | Major capsid protein | 3 |
| 17 | 146812 | 1377 | Tape measure protein | 3 |
| 29 | 130652 | 115 | NKF; TMD | 3 |
| 34 | 165905 | 229 | NKF | 3 |
| 43 | 166702 | 58 | NKF; TMD | 3 |
| 44 | 850 | 68 | NKF | 3 |
| 47 | 167100 | 141 | NKF; DUF3310 | 3 |
| 49 | 164772 | 183 | DnaQ-like DNA pol III subunit | 3 |
| 57 | 84992 | 865 | DNA primase/helicase | 3 |
| 73 | 164822 | 71 | Toxin in toxin/antitoxin system, HicA-like | 3 |

<sup>a</sup> Phamily numbers were recorded from phagesdb.org as of June 3, 2024.
<sup>b</sup> Growth of strains on media supplemented with 10 ng/ml or 100 ng/ml aTc was compared to the same strains plated on media without aTc. For those scored as 0, aTc-dependent reduction in host growth was not observed; those scored as 1 exhibit an aTc-dependent reduction in size colony, those denoted as 2 demonstrate a ~1-3-log difference in growth in the presence of aTc, and those denoted as 3 demonstrate a severe >3-log reduction in growth in the presence of aTc.
<sup>c</sup> NKF, no known function.
<sup>d</sup> TMD, transmembrane domain.

**Figure 1.**
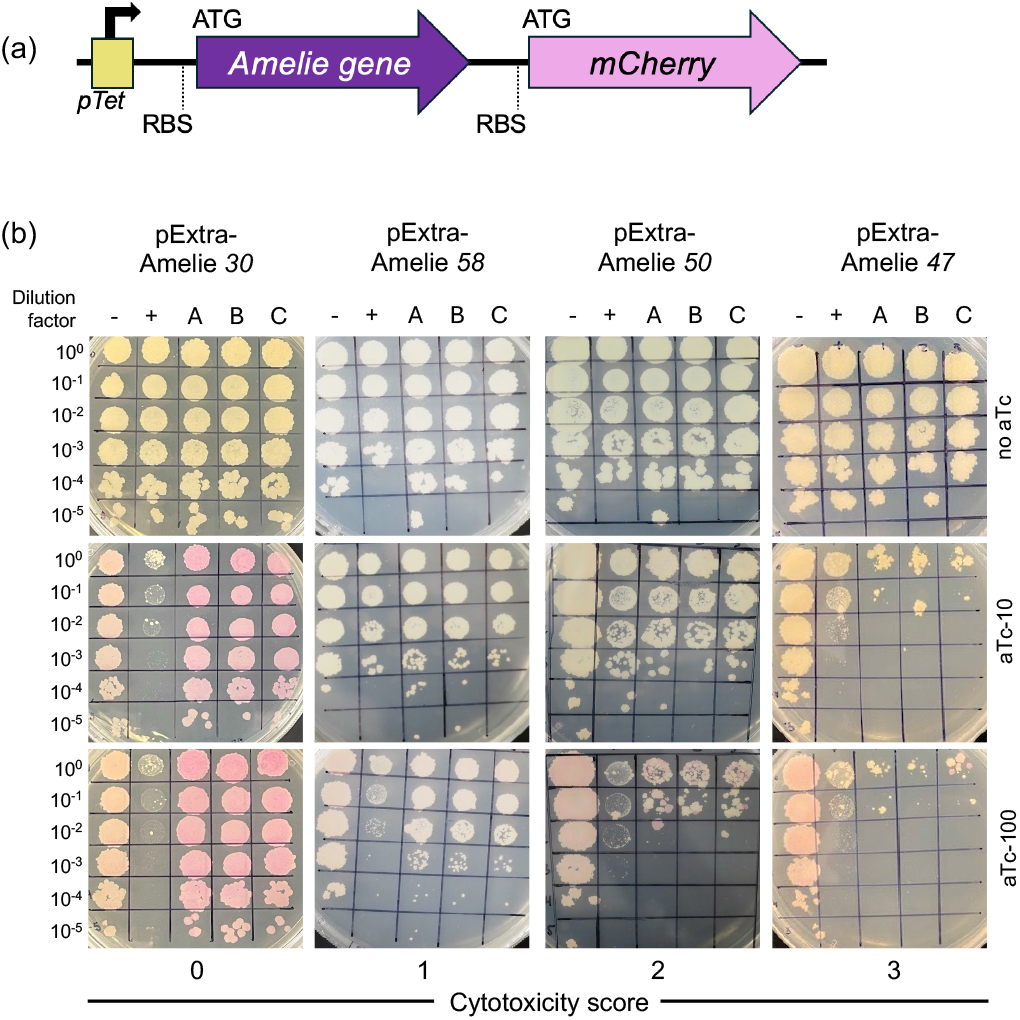
Results of cytotoxicity assays for representative Amelie genes. A) Recombinant pExTra plasmids constructed in this study encode Amelie gene sequences downstream of the *pTet* promoter and upstream of *mcherry*. The two genes in this aTc-inducible operon are transcriptionally linked, with each gene having distinct start and stop codons and ribosomal binding sites (RBS) for translation of the two protein products. B) Results of representative cytotoxicity assays are shown to demonstrate the range of observed growth defects. In each assay, colonies of *M. smegmatis* mc^2^155 transformed with the specified pExTra plasmid were resuspended, serially diluted, and spotted on 7H10 Kan media containing 0, 10, or 100 ng/ml aTc. Triplicate colonies (A, B, C) were tested for each gene alongside a positive control strain (+) transformed with pExTra02 (expressing wildtype Fruitloop *52*) and a negative control strain (−) transformed with pExTra03 (expressing Fruitloop *52 I70S*). Cytotoxicity score (0) is represented by Amelie *30*. Cytotoxicity score (1) is represented by Amelie *58*. Cytotoxicity score (2) is represented by Amelie *50*. Cytotoxicity score (3) is represented by Amelie *47*.

### Cytotoxicity Screening and Phenotype Scoring

To assess cytotoxicity, three colonies of *M. smegmatis* mc^2^155 transformed with individual pExTra-Amelie gene plasmids were resuspended and serially diluted in 7H9 broth then spotted on 7H10 plates supplemented with 10 µg/ml Kanamycin and 0, 10, or 100 ng/ml anhydrotetracycline (aTc; IBA LifeSciences). Each strain was tested in triplicate alongside the pExTra02 positive control plasmid, encoding cytotoxic gene Fruitloop *52*, and the pExTra03 negative control plasmid, encoding a non-toxic mutant allele of Fruitloop *52* (I70S) (Ko and Hatfull 2018; Heller *et al*. 2022). Plates were incubated at 37°C and growth was monitored over 5 days. Two factors were used to evaluate potential cytotoxic effects: reduction in growth with aTc induction and relative growth compared to the control strains.

Cytotoxic effects were classified into four categories based on the extent of growth inhibition: no observable effect on cell viability (score 0), reduced colony size compared to the negative controls, indicating partial toxicity (score 1), moderate toxicity with 1-3 log decrease in viable cell count (score 2), and severe cytotoxicity, causing complete or near complete (>3-log) inhibition of growth (score 3). In cases where no cytotoxicity was observed (score 0), the presence of pink colonies on aTc plates indicated successful mCherry expression driven by the pTet operon, confirming gene induction.

Cytotoxicity for all genes, with the exception of gene *6*, was confirmed by performing two to five independent experiments. Cytotoxicity was assigned when growth inhibition was observed in all independent experiments with strong agreement between triplicate samples within each experiment. For some genes, a slight overall growth reduction was observed on aTc plates, affecting even the pExTra03 negative controls. In such cases, cytotoxicity was only scored if the Amelie samples displayed notable additional growth inhibition compared to the negative controls. When cytotoxicity levels varied between experiments, particularly for score 1 genes, the mildest effect was assigned as the final score.

### Amelie genomic analysis

The Amelie genome map was created using the web-based tool Phamerator (www.phamerator.org) (Cresawn *et al*. 2011).

Reported gene functions are based on those available in the Amelie GenBank record (Accession KX808132). A further round of annotation was performed to assign functions to several more genes by using the Phage Evidence Collection and Annotation Network (PECAAN) (http://pecaan.kbrinsgd.org), which collects information from PhagesDB with BLAST (Russell and Hatfull 2017), HHPRED (PDB_mmCIF70_10_Jan, SCOPe70_2.08, Pfam-A_v35, NCBI_Conserved_Domains(CD)_v3.19) (Gabler *et al*. 2020), NCBI BLAST (Altschul *et al*. 1990), the Conserved Domain Database (Marchler-Bauer *et al*. 2015), DeepTMHMM (Hallgren *et al*. 2022), and TOPCONS (Tsirigos *et al*. 2015) for evidence of gene function. Default parameters were used for all software unless otherwise specified. Final functional predictions (Figure 2, Table 1) were based on HHPRED results with high coverage (>50%), high probability (>90%), and e-values < 1 × 10^−5^. Proteins that did not meet the criteria listed above were classified as hypothetical proteins with no known function (NKF). All predicted proteins were evaluated for common domains such as domains of unknown function (DUF) using the NCBI conserved domain database (listed above). Helix-turn-helix (HTH) DNA binding domains were predicted for Amelie hypothetical proteins gp32 and gp65 based on HHPRED alignment to HTH-domain proteins with high probability (>90%) and confirmed using NPS Helix-turn-Helix predictor (https://npsa-prabi.ibcp.fr/). Gene content comparison between Amelie, Hammy and Waterfoul genomes was performed using the gene content comparison tool on phagesDB (https://phagesdb.org/genecontent/), with phamily designations downloaded from the database on June 3, 2024. Amino acid identities of homologous proteins were calculated using UniProt align (The UniProt Consortium 2023).

**Figure 2.**
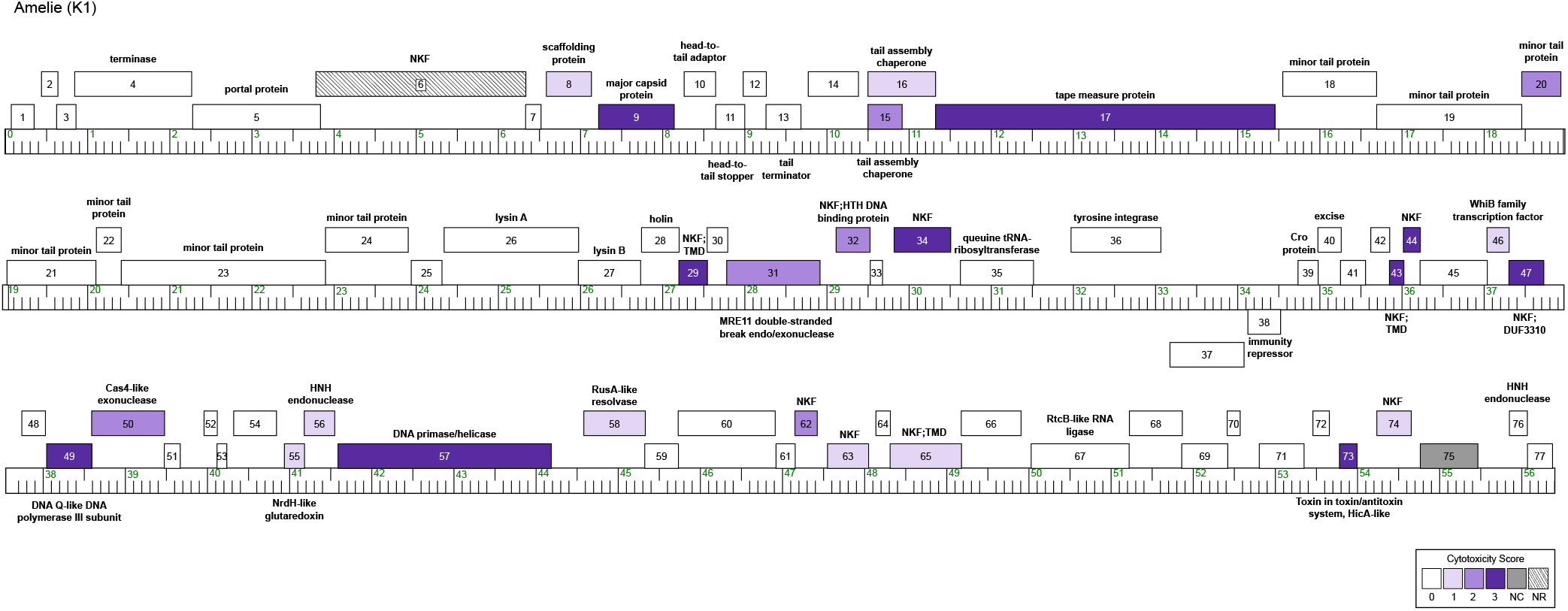
Phage Amelie genome. The Amelie genome is shown as a line with kbp markers and genes represented by boxes—those above the line are transcribed rightwards and those below are transcribed leftwards. Numbers inside the box correspond to gene numbers and predicted functions are indicated above or below each gene. Box shading corresponds to cytotoxicity scoring, with white boxes designating genes found to have no effect on *M. smegmatis* growth (cytotoxicity score *0*), the gray box indicating that the gene was not tested in this study as no clones were obtained (NC), the checkered box indicating a gene for which transformants could not be recovered (NR) and purple representing observed toxicity in our assay. The saturation of purple boxes corresponds to the severity of growth inhibition using the following scores: light purple (score *1*; reduction in colony size; genes *8, 16, 46, 55, 56, 58, 63, 65*, and *74*), medium purple (score *2*; 1–3 log reduction in viability; genes *15, 20, 31, 32, 50*, and *62*), and dark

## Results and Discussion Study Overview

To systematically investigate the effects of Amelie gene overexpression on *M. smegmatis* growth, a gene expression library of Amelie protein coding genes was cloned into the pExtra plasmid under control of the inducible pTet promoter and linked to the *mcherry* fluorescent reporter gene (Fig. 1a). Seventy-six genes were included in this analysis, excluding only gene *75*, as it could not be successfully cloned into the pExtra plasmid despite multiple attempts. Sequence verified gene plasmids were used to transform *M. smegmatis* mc^2^155 for evaluation in a semiquantitative, plate-based cytotoxicity assay.

For each Amelie gene, ten-fold dilution series for three transformed colonies were prepared and spotted on media lacking the aTc inducer, or supplemented with on 10 ng/ml or 100 ng/ml of aTc (Fig. 1b). Control strains expressing wildtype cytotoxic gene Fruitloop *52* or a nontoxic mutant allele (Fruitloop *52*-I70S) were also run alongside the three colonies for each Amelie gene (Ko and Hatfull 2018). Cytotoxic effects of Amelie gene expression were classified into four categories based on the extent of growth inhibition; no observable effect on cell viability (score 0), reduced colony size compared to the negative controls, indicating partial toxicity (score 1), moderate toxicity with 1-3 log decrease in viable cell count (score 2), and severe cytotoxicity, causing complete or near complete (>3-log) inhibition of growth (score 3). Final cytotoxicity scores were assigned based on observations from the higher concentration of inducer. In cases where no cytotoxicity was observed (score 0), the presence of pink colonies on aTc plates indicated successful *mcherry* production driven by the pTet operon, confirming gene induction. For Amelie gene *60*, as was the case for the homologous gene *71* in phage Hammy (Amaya *et al*. 2023), pink color was observed in the absence of aTc, suggesting that this sequence may harbor an internal promoter. Figure 1b shows a representative example of the different levels of cytotoxic activity and the corresponding data and toxicity scores for the entire genome is presented in Supplementary Fig. 1.

### Overexpression of 26 Amelie genes leads to growth inhibition

The overexpression of 50 out of 76 genes screened did not result in an appreciable reduction in bacterial colony growth (Fig 2 and Supplementary Fig. 1). For the majority of genes that had no measurable impact on growth (45 out of 50), *mcherry* expression was visible on at least the aTc-100 plates, confirming expression through the *pTet* operon (Supplementary Fig. 1). It should be pointed out that since Amelie protein levels were not determined in this study we cannot rule out that these gene products may accumulate poorly or not at all in our system.

Thirty four percent of the 76 Amelie protein-coding genes tested were found to reproducibly cause a reduction in *M. smegmatis* growth (Fig. 2, Table 1). This is in line with previous systematic screen for phages Hammy (25% of the genome was cytotoxic, (Amaya *et al*. 2023)), Waterfoul (34%, (Heller *et al*. 2022)), and Girr (28%, (Pollenz *et al*. 2024)). Nine Amelie genes caused effects characterized as mild (cytotoxicity score 1), 6 caused a 1-2 log reduction in colony number (cytotoxicity score 2), and 10 resulted in a >3-log reduction in colony number or almost complete inhibition of growth (cytotoxicity score 3) (Fig. 3a). Gene *43*, which has no known function, encodes the smallest cytotoxic gene product at 58 amino acids while the tape measure protein encodes the largest gene product at 1,377 amino acids (Table 1).

**Figure 3.**
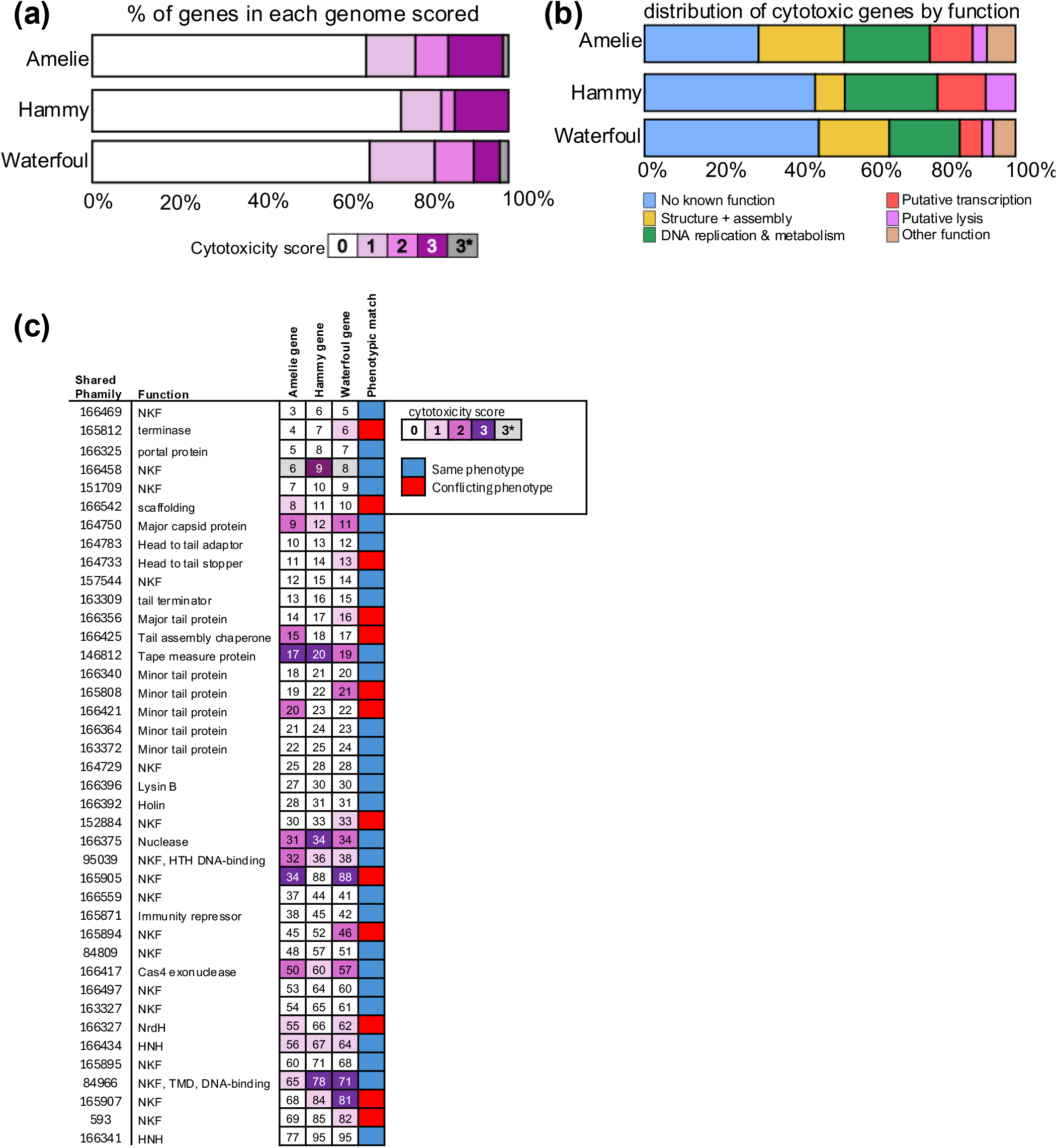
Comparison of Amelie, Hammy and Waterfoul patterns of cytotoxicity. A) The proportion of Amelie (top), Hammy (middle) and Waterfoul (bottom) genes tested in this study or (Heller *et al*. 2022; Amaya *et al*. 2023) that were assigned each score (*0–3*) is represented as a stacked bar chart. Amelie gene 6 as well Waterfoul *8* and *86* were scored as 3* as recovery of pExTra transformants was inhibited by presence of these gene inserts even in the absence of aTc. B) The proportion of Amelie (top), Hammy (middle) and Waterfoul (bottom) cytotoxic genes (score *1*–*3*) that have NKF or that fall into various functional classes is represented as a stacked bar chart with different colors indicating functional class as described in the key. C) Shown is a chart listing the 40 gene phamilies shared by Amelie, Hammy and Waterfoul that were tested in the three studies. Phamily number designations and functions are listed (NKF, no known function; TMD, transmembrane domain) next to representative homologous genes from Amelie, Hammy and Waterfoul, with boxes shaded by cytotoxicity score. A binary indicator of whether homologous genes were both classified as toxic or nontoxic is illustrated by green or red shading, respectively.

Amongst the 26 genes identified as toxic, 16 have predicted functions and although they are scattered throughout the genome, toxic genes are often found near each other. A cluster of six genes involved in structure and assembly is found within the first 20 genes of the genome. Overproduction of the scaffolding protein gp8 and the open reading frame for the longer tail assembly chaperone isoform gp16 caused mild toxicity (score 1), the major capsid protein gp9, the short tail assembly chaperone isoform gp15 and a minor tail protein gp20 caused moderate toxicity (score 2), whereas the tape measure protein gp17 was a more potent growth inhibitor (score 3). Some tape measure proteins contain conserved motifs that may have additional functions beyond just measuring tail length. The tape measure proteins of both Amelie and Waterfoul both contain a short motif that is related to predicted actinobacterial protease proteins (Pedulla *et al*. 2003).

Six gene products that decreased *M. smegmatis* growth are predicted to be involved in functions related to DNA replication or metabolism. These genes included the predicted MRE11 double-strand break exo/endonuclease (gp31), Cas4 family exonuclease (gp50), an HNH endonuclease (gp56), RusA-like resolvase (gp58), and likely replisome components DNA primase/helicase (gp57) and DnaQ-like DNA polymerase III subunit (gp49). Both the DNA primase/helicase (gp57) and DnaQ-like DNA polymerase III subunit (gp49) were highly toxic (Table 1; Supplementary Fig. 1), consistent with replisome function being both essential and sensitive to the stoichiometry of its constituent parts (Reyes-Lamothe *et al*. 2010; Brüning *et al*. 2016; Homiski *et al*. 2021). Although Amelie gp47 could not be assigned a function, it contains a conserved domain of unknown function (DUF3310) that is widespread in phage and bacterial proteomes and was previously implicated in nucleotide kinase activity for the T7 protein gp1.7 (Tran *et al*. 2008, 2012).

Three genes encode for proteins that bind to DNA and likely regulate gene expression. Overproduction of the WhiB transcription factor gp46 caused mild toxicity (Table 1). Additionally, two proteins with predicted helix-turn-helix DNA-binding motifs, gp32 and gp65, caused moderate toxicity (Table 1).

DeepTMHMM predicts that three cytotoxic genes have transmembrane domains (TMDs). In addition to its predicted helix-turn-helix DNA-binding motif near the C-terminus, gp65 is predicted to have 4 transmembrane helices near the N-terminus and a possible helix-turn-helix DNA binding motif near the C-terminus. Unlike the homologous genes in Hammy and Waterfoul, expression of gp65 resulted in only a mild effect on cell growth (Figure 3c). gp29 has a predicted transmembrane helix at its N-terminus and it is likely part of the lysis cassette as it is located downstream of the lysin A and lysin B genes as well as the putative holin gene. Previous studies have shown that phage lysis cassettes often encode multiple holin-like transmembrane domain proteins and that these proteins play a role in controlling the timing of lysis (Catalão *et al*. 2011; Pollenz *et al*. 2022). Although it was the smallest cytotoxic gene product at 58 amino acids, the single transmembrane domain protein gp43 was highly toxic (Table 1); the longest cytotoxic gene product, tape measure protein gp17 also has a single predicted TMD towards its N-terminus. It should be noted that two other predicted membrane proteins, including the putative holin gp28, were not toxic despite bright pink coloration of the cells, suggesting that the proteins were likely produced in our assay (Supplementary Fig. 1). Two other genes that could not be assigned to one of the previous categories were also cytotoxic. Overexpression of the putative NrdH-like glutaredoxin (gp55) caused a mild growth defect in *M. smegmatis*. In contrast, the predicted HicA-like toxin proved to be highly toxic. HicA is a toxin component of the HicAB toxin-antitoxin (TA) system that is widespread in bacterial and archaeal genomes, often associated with mobile genetic elements like prophages (Li *et al*. 2016). Overproduction of HicA in *E. coli* inhibits bacterial growth by cleaving mRNAs and tmRNA in a ribosome-independent manner, thereby reducing the global translation rate and causing cell stasis (Li *et al*. 2016; Chan *et al*. 2023).

Finally, five other genes had no known function and did not have any conserved domains or motifs. Two of them (gp63, gp74) had a mild effect on cell growth, one (gp62) had a moderate effect, and two (gp34, gp44) were highly toxic.

Overall, our screen revealed that overexpressing about one-third of the Amelie genes adversely impacted the growth of *M. smegmatis*. This set of growth-inhibiting genes exhibits diversity in terms of gene size, sequence characteristics, and their impact on cell growth. Despite the artificially elevated protein levels, it is encouraging that 17 out of the 26 genes that encode for cytotoxic proteins elicited either moderate or severe growth defects. In many cases, the effects were evident even at lower inducer concentrations (Supplementary Fig. 1). These potent growth-inhibitory phage products represent potential targets for further analysis in order to elucidate their physiological role in the phage life cycle.

### Conservation of mycobacterial growth inhibition by related phage genes

The next step was to compare the phenotype of each gene phamily encoded by Amelie to the closely related Cluster K phages Waterfoul (Heller *et al*. 2022) and Hammy (Amaya *et al*. 2023). The proteins expressed by the genes within the sequenced genomes of actinobacteriophages are clustered into phamilies (phams) according to similarities in their amino acid sequences (Cresawn *et al*. 2011; Gauthier *et al*. 2022). This categorization based on sequence homology allows for comparative analysis and functional predictions for the Amelie phage genes by leveraging information from related genes across different phages (Fig. 3b). Amelie, Waterfoul and Hammy encode 40 phams in common, 27 shared the same phenotype across all three phages while 13 had conflicting phenotypes (Fig. 3c). Of the 27 shared phams, 19 of them were nontoxic while 8 phams inhibited *M. smegmatis* growth to various degrees. Additionally, although not placed in the same pham, Amelie gp47, Waterfoul gp47, and Hammy gp54 all share a DUF3310 domain on their C-terminus and all three were highly toxic.

Further analysis showed that seven phams present in both Amelie (gp29, gp43, gp44, gp46, gp49, gp57, and gp58) and Hammy (gp32, gp50, gp51, gp53, gp58, gp68, gp69) but absent from Waterfoul were toxic. Moreover, two phams were toxic in Amelie (gp34, gp55) and Waterfoul (gp88, gp62) but not in Hammy (gp88, gp66). Thus, homologous gene products with similar cytotoxic effects represent promising candidates for factors that disrupt critical processes within mycobacterial hosts (Fig. 3b).

There were two structural proteins that were toxic across all three cluster K genomes, the major capsid proteins Amelie gp9, Hammy gp11, and Waterfoul gp12 (>80% identical), and the tape measure proteins Amelie gp17, Hammy gp20, and Waterfoul gp19 (62-71% identical). Minor tail protein Amelie gp20 was moderately toxic but Hammy gp23 and Waterfould gp22, which are all part of the same pham, were not. Amelie gp20 is 71% identical to Hammy gp23 and 64% identical to Waterfoul gp22. It should be noted that a different minor tail protein, Waterfoul gp21, was also moderately toxic but Amelie gp19 and Hammy gp22 were not toxic despite being 75% and 84% identical to it, respectively. Three other structural proteins were toxic in Amelie but the homologs in Hammy and Waterfoul were either not toxic or were not tested (Fig. 3c). Scaffolding protein Amelie gp8 was mildly toxic but Hammy gp11 and Waterfoul gp10 were not, however, these proteins share <19% amino acid similarity to the Amelie scaffolding protein. Moreover, the tail assembly protein Amelie gp15 was moderately toxic while Hammy gp18 and Waterfoul gp17 were not toxic despite sharing >63% similarity. Although numerous structural genes that were toxic lack any known enzymatic activity, they are expressed at high levels during lytic infection (Dedrick *et al*. 2013, 2019a) and their expression may trigger diverse bacterial responses that could affect growth (Zhang *et al*. 2022; Stokar-Avihail *et al*. 2023).

Amelie gp32, a predicted helix-turn-helix DNA binding protein, was moderately toxic whereas Hammy gp36 and Waterfoul gp38 were only mildly toxic. Amelie gp32 is 61% identical to Hammy gp36 and 53% identical to Waterfoul gp38.

Previous studies have shown that overproduction of cII of phage λ, which encodes for a helix-turn-helix motif, was highly toxic although the mechanism of toxicity is not fully understood (Kędzierska *et al*. 2003). The mildly toxic effect of Amelie gp46 is consistent with the phenotype produced by the homologous WhiB-like proteins Hammy gp53 and TM4 gp49 (Rybniker *et al*. 2010). The WhiB-like protein from mycobacteriophage TM4 was previously found to act as an inhibitor of the essential *M. smegmatis* WhiB2 protein, which regulates cell division and septation, by binding to the promoter region of the *whiB2* gene and downregulating its expression (Rybniker *et al*. 2010). Thus, the phage WhiB2 protein acts as a dominant negative inhibitor of the host WhiB2 function, affecting cell growth. It is likely that Amelie gp46 may inhibit cell growth by similarly disrupting normal host gene regulation.

Different phams with putative roles in genome replication were identified as cytotoxic across the three Cluster K phages. These include Amelie gp57 and Hammy gp68 DNA primase/helicase proteins, the Waterfoul sliding clamp subunit gp48, Waterfoul DNA helicase protein gp66, Waterfoul DNA primase/polymerase gp65, and the DnaQ-like DNA pol III subunit proteins Amelie gp49 and Hammy gp58. Amelie and Hammy do not encode genes homologous to the Waterfoul DNA primase/helicase, the sliding clamp or the helicase proteins. The DnaQ gene products in Amelie and Hammy are categorized in the same pham and are highly toxic whereas the Waterfoul gp54 is in a different pham and was not toxic. Recent studies on phage-host dynamics have also observed that phage-encoded replication proteins can elicit diverse cellular defense responses (Stokar-Avihail *et al*. 2023). Further investigation is necessary to determine the mechanism of action by which certain phage replication proteins confer cytotoxicity, whether it be disrupting DNA replication of the host cell or a yet to be identified mechanism.

Three sets of genes encoding nucleases were also toxic in all three phages. Amelie gp56, Hammy gp67, and Waterfoul gp63 and gp64 are HNH endonucleases that cause a mild growth phenotype when overexpressed. HNH endonucleases are a diverse family of nucleases that share limited amino acid identity and include nicking endonucleases, homing endonucleases, and restriction endonucleases (Mehta *et al*. 2004). Cas4 family exonucleases in Amelie (gp50) and Waterfoul (gp57) were moderately toxic whereas that pham in Hammy (gp60) was only mildly toxic. Finally, the MRE11 double-stranded break endo/exonuclease family proteins in Amelie (gp31) and Waterfoul (gp34) were moderately toxic while Hammy gp34 was highly toxic. The MRE11 double-stranded break endo/exonucleases are essential for the efficient and accurate repair of DNA double-strand breaks, ensuring the stability and integrity of the genome (Lammens *et al*. 2011). Although the various nucleases serve specific functions, cytotoxicity may be a consequence of nonspecific nuclease activity upon overexpression (Rao and Black 1988; Bhattacharyya and Rao 1994).

When it comes to the lysis cassettes, the genes encoding lysin B and holin in Amelie, Hammy and Waterfoul share the same phams and none of them had an effect on *M. smegmatis* growth. In contrast, lysin A Hammy gp29 was mildly toxic whereas Amelie gp26 and Waterfoul gp29 were not. *lysA* is 73% identical between Amelie and Waterfoul and belong to the same pham, whereas Hammy gp29 is in a different pham and is only 27% identical to Amelie gp26. Thus far Hammy gp29 is one of a small number of *lysA* genes that have been shown to cause lysis in bacteria in the absence of a holin protein (Payne and Hatfull 2012). All three phages also encode a gene immediately downstream of the holin with a predicted transmembrane domain at its N-terminus. Amelie gp29 and Hammy gp32 are in the same pham whereas Waterfoul gp32 is in a different pham. Regardless, overexpression of these transmembrane proteins resulted in either moderately (Hammy and Waterfoul) or highly (Amelie) toxic inhibition of *M. smegmatis* growth. Phage lysis cassettes frequently encode multiple holin-like transmembrane domain proteins that may play a role in disrupting the cell envelope (Catalão *et al*. 2011; Pollenz *et al*. 2022).

Although several of the genes identified in Amelie to be toxic had homologs in either Hammy or Waterfoul, 8 of the inhibitors identified in this study (gp8, gp15, gp16, gp20, gp62, gp63, gp73, gp74) represent novel gene phams for which cytotoxicity had not been previously identified, thereby expanding the number of known phage-encoded growth inhibitors.

### Patterns of phenotypic conservation across different mycobacteriophage clusters

In many cases proteins are grouped into different phams due to their low amino acid identity despite performing the same function (Cresawn *et al*. 2011; Gauthier *et al*. 2022). Thus, the function of Amelie toxic genes was also compared to that of phage Girr, a Cluster F1 phage for which a similar screen was published recently (Pollenz *et al*. 2024). The first commonality was that several structural genes were toxic in Girr and the Cluster K phages Amelie, Hammy and Waterfoul. Despite being in different phams and having little amino acid sequence similarity (<25%), overexpression of the major capsid protein from the Cluster K phages Amelie (gp9), Hammy (gp12), and Waterfoul (gp11) and the Cluster F phage Girr (gp6) led to a decrease in *M. smegmatis* growth. Likewise, Girr minor tail protein gp19, which does not share similarity to the minor tail proteins of Amelie gp20 and Waterfoul gp21, was also moderately toxic. Although the scaffolding protein of Hammy and Waterfoul were not toxic, Girr scaffolding protein gp5 and Amelie gp8 were both toxic even though they are only 26% identical. The tape measure proteins of Cluster K phages were either moderately (Waterfoul) or highly toxic (Amelie and Hammy) whereas the tape measure protein of phage Girr was not. It should be noted that the tape measure proteins of Cluster K phages have a transmembrane domain whereas the one from phage Girr lacks one and they also share less than 24% amino acid identity.

Additionally, the tail assembly chaperones of Amelie (gp15 and gp16) were toxic whereas they were not in Hammy, Waterfoul, or Girr.

As was the case with Cluster K phages, phage Girr also encodes several toxic genes involved in DNA replication and modification. Several components known to be part of the replisome (Yao and O’Donnell 2010) were toxic when overexpressed in either Girr or Cluster K phages. Girr encodes for DNA helicase/methylase gp64 and ssDNA binding protein gp62 whereas Amelie gp57 and Hammy gp68 encode for a DNA primase/helicase, Waterfoul gp48 is a sliding clamp subunit, Waterfoul gp66 is a DNA helicase, Waterfoul gp65 is a DNA primase/polymerase, and Amelie gp49 and Hammy gp58 encode for the DnaQ-like DNA pol III subunit proteins. These phages also encode for several nucleases that can cut and modify DNA. Girr encodes for DnaQ-like exonuclease gp37 and two HNH endonucleases, gp63 and gp32. Amelie, Hammy and Waterfoul all have HNH endonucleases that are toxic. Expression of Amelie gp56, Hammy gp67, Waterfoul gp64 and Girr gp63 results in mild growth inhibition whereas Waterfoul gp63 and Girr gp32 confer moderate toxicity. It should be noted that Amelie gp77, Hammy gp95, Waterfoul gp95 and Girr gp103 all encode for HNH endonucleases that were not toxic, thus, not all endonucleases that are overexpressed have an effect on *M. smegmatis* growth.

Despite being assigned to different phams, the lysis cassettes of the Cluster K phages are similar to that of Cluster F1 phage Girr. The *lysA* and *lysB* are followed by a putative holin and a gene that encodes for a single transmembrane domain.

Overexpression of lysin A for Girr gp31 and Hammy gp29 resulted in mild cytotoxicity whereas Amelie gp26 and Waterfoul gp29 were nontoxic. Moreover, neither lysin B nor the holin proteins were toxic for any of the four phages. However, the single transmembrane domain protein immediately downstream of the holin gene was highly toxic despite the low sequence conservation between the four proteins. Although Amelie gp29 and Hammy pg32 are members of the same pham they only share 44% amino acid identity. Likewise, Amelie gp29 is only 20% identical to Waterfoul gp32 and 26% identical to Girr gp35. Studies from F1 cluster phage Ms6 show that gp27, which is in the same pham as Girr gp35, is a single transmembrane protein whose expression inhibits *E. coli* growth in liquid media (Catalão *et al*. 2011). Furthermore, deletion of Ms6 gp27 and gp26, a protein with two transmembrane domains, inhibits phage survival. The identification of cytotoxic non-holin transmembrane domain proteins provides further evidence that lysis cassettes might contain multiple genes to trigger and control the degradation of the host’s cell wall at the end of the lytic cycle (Pollenz *et al*. 2022).

### Additional insights

The discussion above does not include the results for Amelie genes *6* and *75*. Despite multiple attempts to transform *M. smegmatis* with two different versions of the pExtra clone encoding for gp6, we were unable to obtain any colonies. This is not surprising as previous work showed that the homologous gene in Waterfoul, gene *8*, also did not produce any colonies when transformed into *M. smegmatis* (Heller *et al*. 2022) while overproduction of Hammy gp9 was highly toxic (Amaya *et al*. 2023). However, a nonsense mutation of Waterfould gp8 allowed for the recovery of transformants. This suggests that leaky protein production of Amelie gp6 in the absence of the aTc inducer is sufficient to be highly toxic to *M. smegmatis* (designated as 3* in Fig. 3a), bringing the total number of toxic genes to 26. Moreover, despite multiple attempts Amelie gp75 could not be cloned into the pExtra plasmid. Hammy gp92, which is 69% identical to Amelie gp75, was found to be moderately toxic when overexpressed (Amaya *et al*. 2023), suggesting that there might be another possible cytotoxic gene encoded by Amelie.

Gene content comparisons show that Amelie is 67.3% similar to Hammy, 49.6% similar to Waterfoul, and 4.6% similar to Girr. Despite these differences one pattern that emerged is that mycobacteriophages are rich in cytotoxic genes. 34% of Amelie’s genome encodes for proteins that are cytotoxic to *M. smegmatis*, which is in line with previous reports demonstrating that anywhere from 25% to one-third of phage genes tested inhibited *M. smegmatis* growth to various degrees (Heller *et al*. 2022; Amaya *et al*. 2023; Pollenz *et al*. 2024). While it is unlikely that these cytotoxic genes are actively used by the phages to arrest host growth during infection, these findings demonstrate that phage proteins can interact with and potentially inactivate or alter the function of host proteins essential for bacterial growth and survival (Ko and Hatfull 2020).

Another pattern that emerged is that cytotoxic genes are often clustered with other cytotoxic genes (Fig. 2), suggesting that they may work together through various stages of the phage life cycle. For phage Amelie, there are five clusters that can be identified (Fig. 2). Amelie genes *15*-*17* encode for structural proteins, cluster 2 is found between the holin and tyrosine integrase and it includes a holin-like protein with a predicted transmembrane domain at its N-terminus (gp29), a nuclease (gp31), and two NKF genes (gp32, and gp34). Cluster 3 (genes *43, 44, 46, 47, 49*, and *50*) and cluster 4 (genes *55-58*) encode for genes predicted to function in DNA replication, DNA metabolism, and transcription. Interestingly, they are found downstream of cro (gene *39*) and are likely expressed early in the lytic life cycle (Ko and Hatfull 2018), making them good candidates for proteins involved in the takeover of critical cellular processes. Finally, cluster 5 includes genes *62, 63*, and *65*, all of which are NKF. Amelie has a similar genomic architecture and 59.6% gene content identity with cluster K2 phage ZoeJ, for which transcriptomic data show that several of these cytotoxic genes are co-expressed throughout the phage life cycle (Dedrick *et al*. 2019a).

In summary, while several toxic genes identified in Amelie had homologs in Cluster K phages Hammy or Waterfoul, our study reveals eight novel gene phams (gp8, gp15, gp16, gp20, gp62, gp63, gp73, gp74) that were previously unknown to exhibit cytotoxic properties, thereby broadening the repertoire of known phage-encoded growth inhibitors. This highlights that comparisons between similar phage genomes can uncover differences in cytotoxicity even among closely related genes, presenting opportunities for in-depth downstream analyses to understand the factors behind these varied outcomes. Despite the vast number of known mycobacteriophage gene phamilies, only a limited subset has been experimentally investigated to date (Rybniker *et al*. 2008, 2010, 2011; Ko and Hatfull 2018, 2020; Heller *et al*. 2022; Pollenz *et al*. 2022; Amaya *et al*. 2023), advocating for further genome-wide screens of phage genomes in order to identify host vulnerabilities well-suited for drug development as well as learn more about phage biology and evolution.

## Data Availability

All plasmids and plasmid sequences reported in this study are available upon request. The authors affirm that all data necessary for confirming the conclusions of this article are represented fully within the article and its tables and figures.

Extended data, including plasmid Sanger sequencing data and confirmatory cytotoxicity assay data can be found at the SEA-GENES project open access database GenesDB (https://genesdb.org). To access these data on genesDB.org, any user can register for a free account, and once logged in to this account, navigate from the home page to Cluster K1 phage Amelie. Supplemental material is available online.

## Acknowledgements

This work was conducted as part of the HHMI-supported Science Education Alliance GENES (Gene-function Exploration by a Network of Emerging Scientists) project. We thank members of the Science Education Alliance for their research support including Viknesh Sivanathan, Graham Hatfull, Deborah Jacobs-Sera, Steven Cresawn, and Dan Russell. We thank New England Biolabs (NEB) and Integrated DNA Technologies (IDT) for providing reagent support. We would also like to thank the Biology Department and the College of Arts and Sciences at La Sierra University for supporting the SEA programs on campus.

## Notes

https://genesdb.org/

### Competing Interest Statement

The authors have declared no competing interest.

