## Supplemental Figures and Tables for "Genome-wide screen overexpressing mycobacteriophage Amelie genes identifies multiple inhibitors of mycobacterial growth"

#### Gene *1*; Score *0*

Images taken after 5 days at 37 ºC

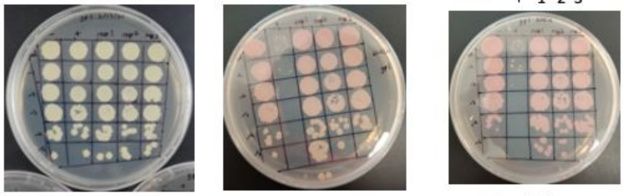


0 ng/ml aTc | 10 ng/ml aTc | 100 ng/ml aTc

| Lane | Gene ID | Plasmid name | Gene name | Toxic/Non-toxic | Colony color on 100 ng/ml aTc plate* |
|---|---|---|---|---|---|
| - Non-toxic control | -- | pExTra03 | Fruitloop *52* mutant | Non-toxic | ++ |
| + Toxic control | -- | pExTra02 | Fruitloop *52* | Toxic | - |
| 1 | | pExTra-Amelie 01 | Amelie 01 replicate 1 | Non-toxic | ++ |
| 2 | | pExTra-Amelie 01 | Amelie 01 replicate 2 | Non-toxic | ++ |
| 3 | | pExTra-Amelie 01 | Amelie 01 replicate 3 | Non-toxic | ++ |

*Key: NG (no growth) - (no pink color) +(faint pink color) ++(obvious pink color) +++ (dark pink color)

#### Gene *2*; Score *0*

Images taken after 5 days at 37 ºC

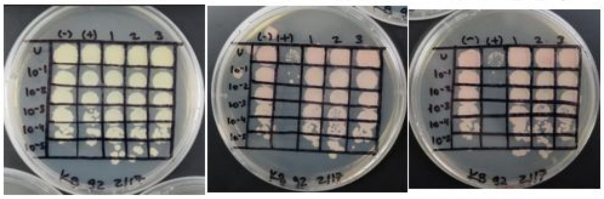


0 ng/ml aTc | 10 ng/ml aTc | 100 ng/ml aTc

| Lane | Gene ID | Plasmid name | Gene name | Toxic/Non-toxic | Colony color on 100 ng/ml aTc plate* |
|---|---|---|---|---|---|
| - Non-toxic control | -- | pExTra03 | Fruitloop *52* mutant | Non-toxic | + |
| + Toxic control | -- | pExTra02 | Fruitloop *52* | Toxic | - |
| 1 | -- | pExTra_2 | Amelie 2 replicate 1 | Non-toxic | + |
| 2 | -- | pExTra_2 | Amelie 2 replicate 2 | Non-toxic | + |
| 3 | -- | pExTra_2 | Amelie 2 replicate 3 | Non-toxic | + |

*Key: NG (no growth) - (no pink color) +(faint pink color) ++(obvious pink color) +++

#### Gene *3*; Score *0*

Images taken after 4 days at 37 ºC

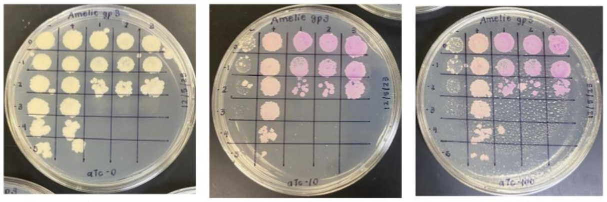


0 ng/ml aTc | 10 ng/ml aTc | 100 ng/ml aTc

| Lane | Gene ID | Plasmid name | Gene name | Toxic/Non-toxic | Colony color on 100 ng/ml aTc plate* |
|---|---|---|---|---|---|
| - Non-toxic control | -- | pExTra03 | Fruitloop *52* mutant | Non-toxic | ++ |
| + Toxic control | -- | pExTra02 | Fruitloop *52* | Toxic | - |
| 1 | | pExTra-Amelie3 | Amelie *3* replicate 1 | Non-toxic | +++ |
| 2 | | pExTra-Amelie3 | Amelie *3* replicate 2 | Non-toxic | +++ |
| 3 | | pExTra-Amelie3 | Amelie *3* replicate 3 | Non-toxic | +++ |

*Key: NG (no growth) - (no pink color) +(faint pink color) ++(obvious pink color) +++ (dark pink color)

#### Gene *4*; Score *0*

Images taken after 4 days at 37 ºC

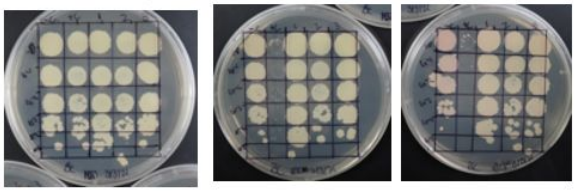


0 ng/ml aTc | 10 ng/ml aTc | 100 ng/ml aTc

| Lane | Gene ID | Plasmid name | Gene name | Toxic/Non-toxic | Colony color on 100 ng/ml aTc plate* |
|---|---|---|---|---|---|
| - Non-toxic control | | pExTra03 | Fruitloop *52* mutant | Non-toxic | ++ |
| + Toxic control | | pExTra02 | Fruitloop *52* | Toxic | - |
| 1 | | pExTra_4 | Amelie 4 replicate 1 | Non-toxic | + |
| 2 | | pExTra_4 | Amelie 4 replicate 2 | Non-toxic | + |
| 3 | | pExTra_4 | Amelie 4 replicate 3 | Non-toxic | + |

*Key: NG (no growth) - (no pink color) +(faint pink color) ++(obvious pink color) +++ (dark pink color)

#### Gene *5*; Score *0*

Images taken after 5 days at 37 ºC

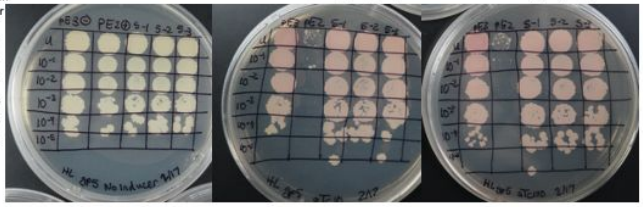


0 ng/ml aTc | 10 ng/ml aTc | 100 ng/ml aTc

| Lane | Gene ID | Plasmid name | Gene name | Toxic/Non-toxic | Colony color on 100 ng/ml aTc plate* |
|---|---|---|---|---|---|
| - Non-toxic control | -- | pExTra03 | Fruitloop *52* mutant | Non-toxic | + |
| + Toxic control | -- | pExTra02 | Fruitloop *52* | Toxic | - |
| 1 | | pExtra_5 | Amelie 5 replicate 1 | Non-toxic | ++ |
| 2 | | pExtra_5 | Amelie 5 replicate 2 | Non-toxic | ++ |
| 3 | | pExtra_5 | Amelie 5 replicate 3 | Non-toxic | ++ |

*Key: NG (no growth) - (no pink color) +(faint pink color) ++(obvious pink color) +++ (dark pink color)

#### Gene *7*; Score *0*

Images taken after 5 days at 37°C

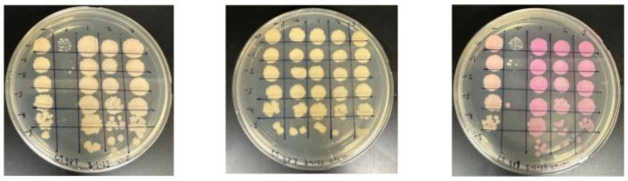


0 ng/ml aTc | 10 ng/ml aTc | 100 ng/ml aTc

| Lane | Gene ID | Plasmid Name | Gene Name | Toxic/Non-Toxic | Colony Color on 100 ng/mL aTc Plate |
|---|---|---|---|---|---|
| (-) Non-Toxic Control | ----- | pExTra_03 | Fruitloop *52* Mutant | Non-Toxic | + |
| (+) Toxic Control | ----- | pExTra_02 | Fruitloop *52* | Toxic | - |
| 1 | ----- | pExTra-Amelie_7 | Amelie 7 Replicate 1 | Non-Toxic | +++ |
| 2 | ----- | pExTra-Amelie_7 | Amelie 7 Replicate 2 | Non-Toxic | +++ |
| 3 | ----- | pExTra-Amelie_7 | Amelie 7 Replicate 3 | Non-Toxic | +++ |

*Key: NG (No Growth) - (no pink color) + (faint pink color) ++ (obvious pink color) +++ (dark pink color)

#### Gene *8*; Score *1*

Images taken after 5 days at 37 ºC

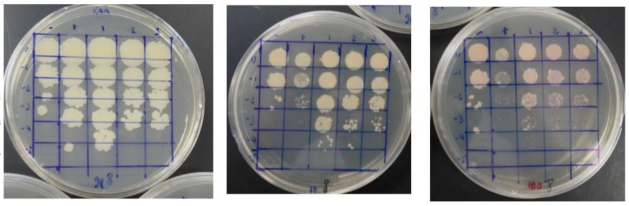


0 ng/ml aTc | 10 ng/ml aTc | 100 ng/ml aTc

| Lane | Gene ID | Plasmid Name | Gene name | Toxic/Non-toxic | Colony color on 100 ng/ml aTc plate* |
|---|---|---|---|---|---|
| - Non-toxic control | -- | pExTra03 | Fruitloop *52* mutant | Non-toxic | + |
| + Toxic control | -- | pExTra02 | Fruitloop *52* | Toxic | - |
| 1 | -- | pExTra-Amelie 8 | Amelie *8* replicate 1 | Toxic | ++ |
| 2 | -- | pExTra-Amelie 8 | Amelie *8* replicate 2 | Toxic | ++ |
| 3 | -- | pExTra-Amelie 8 | Amelie *8* replicate 3 | Toxic | ++ |

*Key: NG (no growth) - (no pink color) +(faint pink color) ++(obvious pink color) +++ (dark pink color)

#### Gene *9*; Score *3*

Images taken after 4 days at 37 ºC

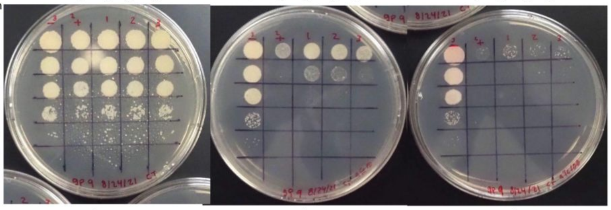


0 ng/ml aTc | 10 ng/ml aTc | 100 ng/ml aTc

| Lane | Gene ID | Plasmid name | Gene name | Toxic/Non-toxic | Colony color on 100 ng/ml aTc plate* |
|---|---|---|---|---|---|
| - Non-toxic control | -- | pExTra03 | Fruitloop *52* mutant | Non-toxic | + |
| + Toxic control | -- | pExTra02 | Fruitloop *52* | Toxic | - |
| 1 | | pExTra-Amelie*09* | Amelie *09* replicate 1 | Toxic | ++ |
| 2 | | pExTra-Amelie09 | Amelie *09* replicate 2 | Toxic | ++ |
| 3 | | pExTra-Amelie09 | Amelie *09* replicate 3 | Toxic | ++ |

*Key: NG (no growth) - (no pink color) +(faint pink color) ++(obvious pink color) +++ (dark pink color)

#### Gene *10*; Score *0*

Images taken after 5 days at 37 ºC

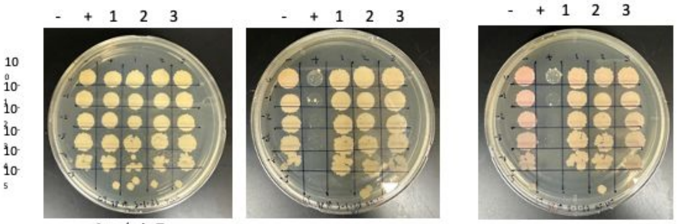


0 ng/ml aTc 10 ng/ml aTc 100 ng/ml aTc

| Lane | Gene ID | Plasmid name | Gene name | Toxic/Non-toxic | Colony color on 100 ng/ml aTc plate* |
|---|---|---|---|---|---|
| - Non-toxic control | -- | pExTra03 | Fruitloop *52* mutant | Non-toxic | ++ |
| + Toxic control | -- | pExTra02 | Fruitloop *52* | Toxic | - |
| 1 | -- | pExTra_10 | Amelie *10* replicate 1 | Non-toxic | ++ |
| 2 | -- | pExTra_10 | Amelie *10* replicate 2 | Non-toxic | ++ |
| 3 | -- | pExTra_10 | Amelie *10* replicate 3 | Non-toxic | ++ |

*Key: NG (no growth) - (no pink color) +(faint pink color) ++(obvious pink color) +++ (dark pink color)

#### Gene *11*; Score *0*

Images taken after 5 days at 37 ºC

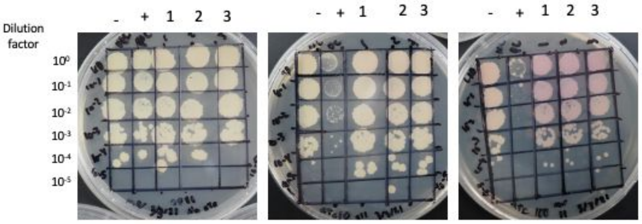


0 ng/ml aTc 10 ng/ml aTc 100 ng/ml aTc

| Lane | Gene ID | Plasmid name | Gene name | Toxic/Non-toxic | Colony color on 100 ng/ml aTc plate* |
|---|---|---|---|---|---|
| - Non-toxic control | -- | pExTra03 | Fruitloop *52* mutant | Non-toxic | + |
| + Toxic control | -- | pExTra02 | Fruitloop *52* | Toxic | - |
| 1 | -- | pExTra_11 | Amelie *11* replicate 1 | Non-toxic | ++ |
| 2 | -- | pExTra_11 | Amelie *11* replicate 2 | Non-toxic | ++ |
| 3 | -- | pExTra_11 | Amelie *11* replicate 3 | Non-toxic | ++ |

*Key: NG (no growth) - (no pink color) +(faint pink color) ++(obvious pink color) +++ (dark pink color)

#### Gene *12*; Score *0*

Images taken after 5 days at 37 ºC

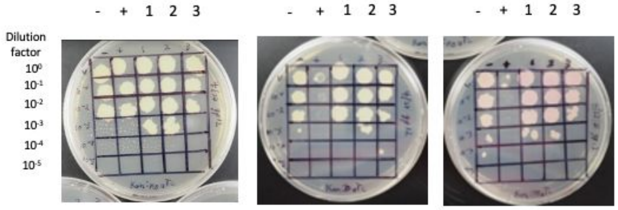


0 ng/ml aTc 10 ng/ml aTc 100 ng/ml aTc

| Lane | Gene ID | Plasmid name | Gene name | Toxic/Non-toxic | Colony color on 100 ng/ml aTc plate* |
|---|---|---|---|---|---|
| - Non-toxic control | -- | pExTra03 | Fruitloop *52* mutant | Non-toxic | + |
| + Toxic control | -- | pExTra02 | Fruitloop *52* | Toxic | - |
| 1 | -- | pExTra_12 | Amelie *12* replicate 1 | Non-toxic | + |
| 2 | -- | pExTra_12 | Amelie *12* replicate 2 | Non-toxic | + |
| 3 | -- | pExTra_12 | Amelie *12* replicate 3 | Non-toxic | + |

*Key: NG (no growth) - (no pink color) +(faint pink color) ++(obvious pink color) +++ (dark pink color)

#### Gene *13*; Score *0*

Images taken after 4 days at 37 ºC

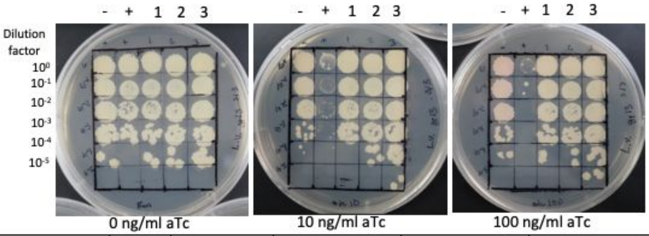


0 ng/ml aTc 10 ng/ml aTc 100 ng/ml aTc

| Lane | Gene ID | Plasmid name | Gene name | Toxic/Non-toxic | Colony color on 100 ng/ml aTc plate* |
|---|---|---|---|---|---|
| - Non-toxic control | -- | pExTra03 | Fruitloop *52* mutant | Non-toxic | - |
| + Toxic control | -- | pExTra02 | Fruitloop *52* | Toxic | NG |
| 1 | -- | pExTra__13 | Amelie 13 replicate 1 | Non-toxic | - |
| 2 | -- | pExTra__13 | Amelie 13 replicate 1 | Non-toxic | - |
| 3 | -- | pExTra__13 | Amelie 13 replicate 1 | Non-toxic | - |

*Key: NG (no growth) - (no pink color) +(faint pink color) ++(obvious pink color) +++

#### Gene *14*; Score *0*

Images taken after 4 days at 37 ºC

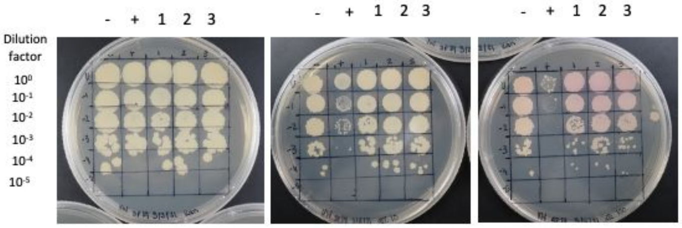


0 ng/ml aTc 10 ng/ml aTc 100 ng/ml aTc

| Lane | Gene ID | Plasmid name | Gene name | Toxic/Non-toxic | Colony color on 100 ng/ml aTc plate* |
|---|---|---|---|---|---|
| - Non-toxic control | -- | pExTra03 | Fruitloop *52* mutant | Non-toxic | + |
| + Toxic control | -- | pExTra02 | Fruitloop *52* | Toxic | - |
| 1 | -- | pExTra-Amelie 14 | Amelie 14 replicate 1 | Non-toxic | ++ |
| 2 | -- | pExTra-Amelie 14 | Amelie 14 replicate 2 | Non-toxic | ++ |
| 3 | -- | pExTra-Amelie 14 | Amelie 14 replicate 3 | Non-toxic | ++ |

*Key: NG (no growth) - (no pink color) +(faint pink color)++(obvious pink color) +++ (dark pink color)

#### Gene *15*; Score *2*

Images taken after 4 days at 37 ºC

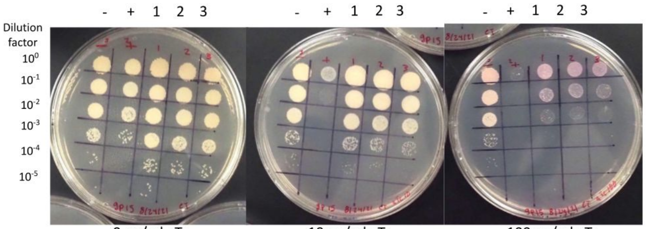


0 ng/ml aTc 10 ng/ml aTc 100 ng/ml aTc

| Lane | Gene ID | Plasmid name | Gene name | Toxic/Non-toxic | Colony color on 100 ng/ml aTc plate* |
|---|---|---|---|---|---|
| - Non-toxic control | -- | pExTra03 | Fruitloop *52* mutant | Non-toxic | + |
| + Toxic control | -- | pExTra02 | Fruitloop *52* | Toxic | - |
| 1 | | pExTra-Amelie15 | Amelie 15 replicate 1 | Toxic | +++ |
| 2 | | pExTra-Amelie15 | Amelie 15 replicate 2 | Toxic | +++ |
| 3 | | pExTra-Amelie15 | Amelie 15 replicate 3 | Toxic | +++ |

*Key: NG (no growth) - (no pink color) +(faint pink color) ++(obvious pink color) +++ (dark pink color)

#### Gene *16*; Score *1*

Images taken after 4 days at 37 ºC

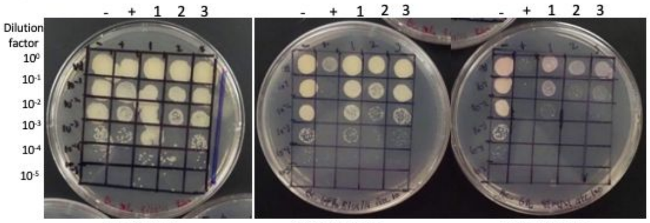


0 ng/ml aTc 10 ng/ml aTc 100 ng/ml aTc

| Lane | Gene ID | Plasmid name | Gene name | Toxic/Non-toxic | Colony color on 100 ng/ml aTc plate* |
|---|---|---|---|---|---|
| - Non-toxic control | -- | pExTra03 | Fruitloop *52* mutant | Non-toxic | + |
| + Toxic control | -- | pExTra02 | Fruitloop *52* | Toxic | - |
| 1 | | pExTra-Amelie16 | Amelie 16 replicate 1 | Toxic | ++ |
| 2 | | pExTra-Amelie16 | Amelie 16 replicate 2 | Toxic | ++ |
| 3 | | pExTra-Amelie16 | Amelie 16 replicate 3 | Toxic | ++ |

*Key: NG (no growth) - (no pink color) +(faint pink color) ++(obvious pink color) +++ (dark pink color)

#### Gene *17*; Score *3*

Images taken after 5 days at 37 ºC

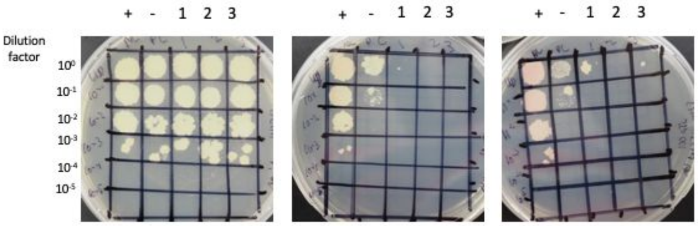


0 ng/ml aTc 10 ng/ml aTc 100 ng/ml aTc

| Lane | Gene ID | Plasmid name | Gene name | Toxic/Non-toxic | Colony color on 100 ng/ml aTc plate* |
|---|---|---|---|---|---|
| - Non-toxic control | -- | pExTra03 | Fruitloop *52* mutant | Non-toxic | + |
| + Toxic control | -- | pExTra02 | Fruitloop *52* | Toxic | - |
| 1 | -- | pExTra-Amelie 17 | Amelie *17* replicate 1 | Toxic | - |
| 2 | -- | pExTra-Amelie 17 | Amelie *17* replicate 2 | Toxic | NG |
| 3 | -- | pExTra-Amelie 17 | Amelie *17* replicate 3 | Toxic | NG |

*Key: NG (no growth) - (no pink color) +(faint pink color) ++(obvious pink color) +++ (dark pink color)

#### Gene *18*; Score *0*

Images taken after 5 days at 37 ºC

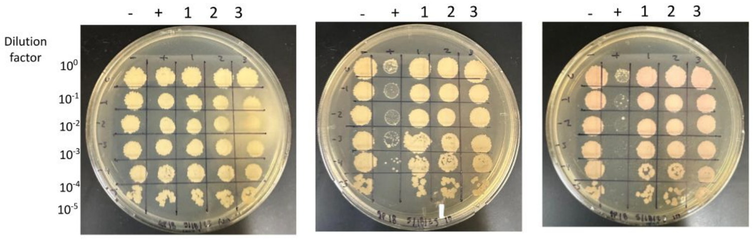


0 ng/ml aTc 10 ng/ml aTc 100 ng/ml aTc

| Lane | Gene ID | Plasmid name | Gene name | Toxic/Non-toxic | Colony color on 100 ng/ml aTc plate* |
|---|---|---|---|---|---|
| - Non-toxic control | -- | pExTra03 | Fruitloop *52* mutant | Non-toxic | + |
| + Toxic control | -- | pExTra02 | Fruitloop *52* | Toxic | - |
| 1 | -- | pExTra-Amelie 18 | Amelie 18 replicate 1 | Non-toxic | ++ |
| 2 | -- | pExTra-Amelie 18 | Amelie 18 replicate 2 | Non-toxic | ++ |
| 3 | -- | pExTra-Amelie 18 | Amelie 18 replicate 3 | Non-toxic | ++ |

*Key: NG (no growth) - (no pink color) +(faint pink color) ++(obvious pink color) +++ (dark pink color)

#### Gene *19*; Score 0

Images taken after 5 days at 37 ºC

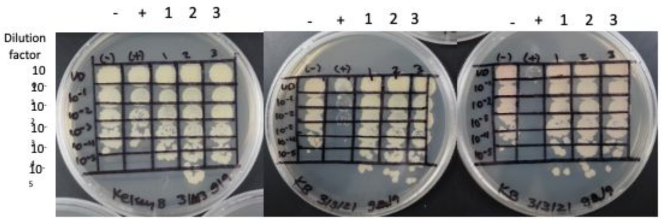


0 ng/ml aTc 10 ng/ml aTc 100 ng/ml aTc

| Lane | Gene ID | Plasmid name | Gene name | Toxic/Non-toxic | Colony color on 100 ng/ml aTc plate* |
|---|---|---|---|---|---|
| - Non-toxic control | -- | pExTra03 | Fruitloop *52* mutant | Non-toxic | + |
| + Toxic control | -- | pExTra02 | Fruitloop *52* | Toxic | - |
| 1 | -- | pExTra_19 | Amelie 19 replicate 1 | Non-toxic | + |
| 2 | -- | pExTra_19 | Amelie 19 replicate 2 | Non-toxic | + |
| 3 | -- | pExTra_19 | Amelie 19 replicate 3 | Non-toxic | + |

*Key: NG (no growth) - (no pink color) +(faint pink color) ++(obvious pink color) +++ (dark pink color)

#### Gene *20*; Score 2

Images taken after 4 days at 37 ºC

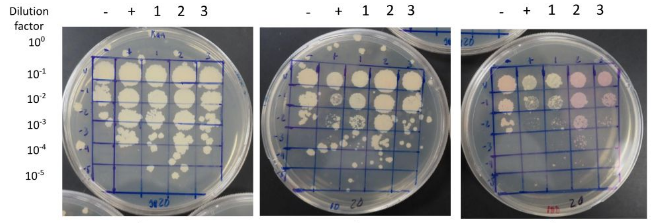


0 ng/ml aTc 10 ng/ml aTc 100 ng/ml aTc

| Lane | Gene ID | Plasmid name | Gene name | Toxic/Non-toxic | Colony color on 100 ng/ml aTc plate* |
|---|---|---|---|---|---|
| - Non-toxic control | -- | pExTra03 | Fruitloop *52* mutant | Non-toxic | + |
| + Toxic control | -- | pExTra02 | Fruitloop *52* | Toxic | - |
| 1 | | pExTra-Amelie20 | Amelie 20 replicate 1 | Toxic | ++ |
| 2 | | pExTra-Amelie20 | Amelie 20 replicate 2 | Toxic | ++ |
| 3 | | pExTra-Amelie20 | Amelie 20 replicate 3 | Toxic | ++ |

*Key: NG (no growth) - (no pink color) +(faint pink color) ++(obvious pink color) +++ (dark pink color)

#### Gene *21*; Score *0*

Images taken after 5 days at 37 ºC

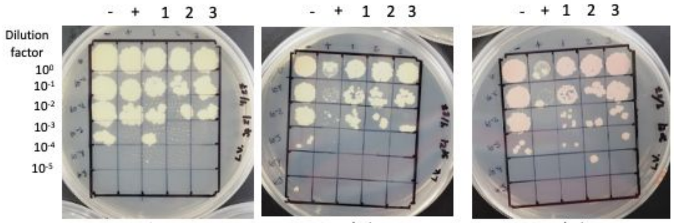


0 ng/ml aTc 10 ng/ml aTc 100 ng/ml aTc

| Lane | Gene ID | Plasmid name | Gene name | Toxic/Non-toxic | Colony color on 100 ng/ml aTc plate* |
|---|---|---|---|---|---|
| - Non-toxic control | -- | pExTra03 | Fruitloop *52* mutant | Non-toxic | + |
| + Toxic control | -- | pExTra02 | Fruitloop *52* | Toxic | - |
| 1 | -- | pExTra__Amelie21 | Amelie 21 replicate 1 | Non-toxic | ++ |
| 2 | -- | pExTra__Amelie21 | Amelie 21 replicate 2 | Non-toxic | ++ |
| 3 | -- | pExTra__Amelie21 | Amelie 21 replicate 3 | Non-toxic | ++ |

*Key: NG (no growth) - (no pink color) +(faint pink color) ++(obvious pink color) +++ (dark pink color)

#### Gene *22*; Score *0*

Images taken after 5 days at 37 ºC

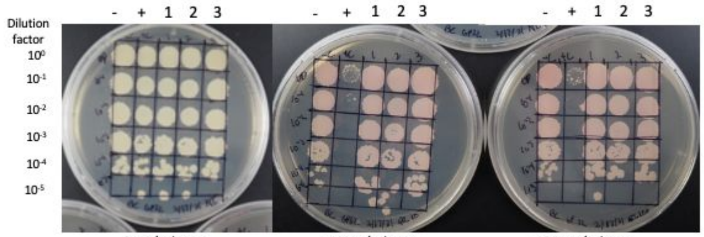


0 ng/ml aTc 10 ng/ml aTc 100 ng/ml aTc

| Lane | Gene ID | Plasmid name | Gene name | Toxic/Non-toxic | Colony color on 100 ng/ml aTc plate* |
|---|---|---|---|---|---|
| - Non-toxic control | | pExTra03 | Fruitloop *52* mutant | Non-toxic | ++ |
| + Toxic control | | pExTra02 | Fruitloop *52* | Toxic | - |
| 1 | | pExTra_22 | Amelie *22* replicate 1 | Non-toxic | ++ |
| 2 | | pExTra_22 | Amelie *22* replicate 2 | Non-toxic | ++ |
| 3 | | pExTra_22 | Amelie *22* replicate 3 | Non-toxic | ++ |

*Key: NG (no growth) - (no pink color) +(faint pink color) ++(obvious pink color) +++ (dark pink color)

#### Gene *23*; Score *0*

Images taken after 5 days at 37 ºC

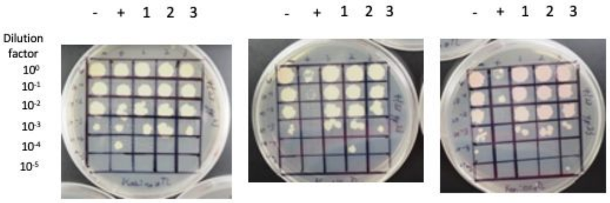


0 ng/ml aTc 10 ng/ml aTc 100 ng/ml aTc

| Lane | Gene ID | Plasmid name | Gene name | Toxic/Non-toxic | Colony color on 100 ng/ml aTc plate* |
|---|---|---|---|---|---|
| - Non-toxic control | -- | pExTra03 | Fruitloop *52* mutant | Non-toxic | + |
| + Toxic control | -- | pExTra02 | Fruitloop *52* | Toxic | - |
| 1 | -- | pExTra_23 | Amelie *23* replicate 1 | Non-toxic | + |
| 2 | -- | pExTra_23 | Amelie *23* replicate 2 | Non-toxic | + |
| 3 | -- | pExTra_23 | Amelie *23* replicate 3 | Non-toxic | + |

*Key: NG (no growth) - (no pink color) +(faint pink color) ++(obvious pink color) +++ (dark pink color)

#### Gene *24*; Score *0*

Images taken after 5 days at 37 ºC

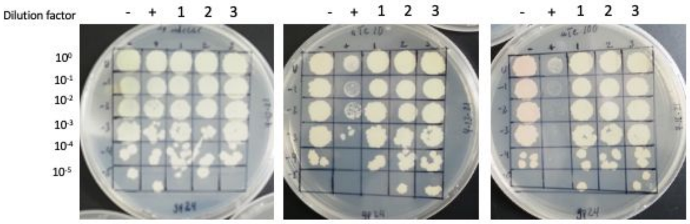


0 ng/ml aTc 10 ng/ml aTc 100 ng/ml aTc

| Lane | Gene ID | Plasmid name | Gene name | Toxic/Non-toxic | Colony color on 100 ng/ml aTc plate* |
|---|---|---|---|---|---|
| - Non-toxic control | -- | pExTra03 | Fruitloop *52* mutant | Non-toxic | + |
| + Toxic control | -- | pExTra02 | Fruitloop *52* | Toxic | - |
| 1 | | pExTra- Amelie 24 | Amelie 24 replicate 1 | Non-toxic | - |
| 2 | | pExTra- Amelie 24 | Amelie 24 replicate 2 | Non-toxic | - |
| 3 | | pExTra- Amelie 24 | Amelie 24 replicate 3 | Non-toxic | - |

*Key: NG (no growth) - (no pink color) +(faint pink color) ++(obvious pink color) +++ (dark pink color)

#### Gene *25*; Score *0*

Images taken after 5 days at 37°C

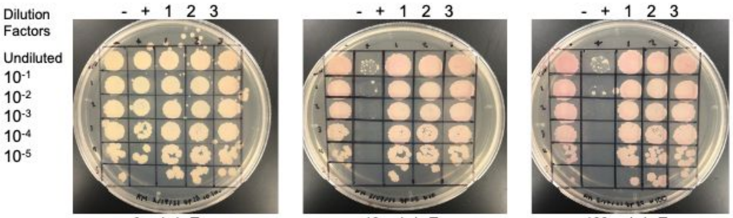


0 ng/ml aTc 10 ng/ml aTc 100 ng/ml aTc

| Lane | Gene ID | Plasmid Name | Gene Name | Toxic/Non-Toxic | Colony Color on 100 ng/mL aTc Plate |
|---|---|---|---|---|---|
| (-) Non-Toxic Control | ----- | pExTra_03 | Fruitloop *52* Mutant | Non-Toxic | ++ |
| (+) Toxic Control | ----- | pExTra_02 | Fruitloop *52* | Toxic | - |
| 1 | ----- | pExTra_25 | Amelie 25 Replicate 1 | Non-Toxic | ++ |
| 2 | ----- | pExTra_25 | Amelie 25 Replicate 2 | Non-Toxic | ++ |
| 3 | ----- | pExTra_25 | Amelie 25 Replicate 3 | Non-Toxic | ++ |

*Key: NG (No Growth) - (no pink color) + (faint pink color) ++ (obvious pink color) +++ (dark pink color)

#### Gene *26*; Score *0*

Images taken after 5 days at 37 ºC

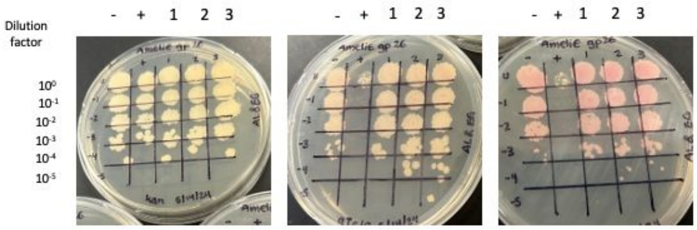


0 ng/ml aTc** | 10 ng/ml aTc | 100 ng/ml aTc

| Lane | Gene ID | Plasmid name | Gene name | Toxic/Non-toxic | Colony color on 100 ng/ml aTc plate* |
|---|---|---|---|---|---|
| - Non-toxic control | -- | pExTra03 | Fruitloop *52* mutant | Non-toxic | + |
| + Toxic control | -- | pExTra02 | Fruitloop *52* | Toxic | + |
| 1 | -- | pExTra_Amelie26 | Amelie *26* replicate 1 | Non-toxic | + |
| 2 | -- | pExTra_Amelie26 | Amelie *26* replicate 2 | Non-toxic | + |
| 3 | -- | pExTra_Amelie26 | Amelie *26* replicate 3 | Non-toxic | + |

*Key: NG (no growth) - (no pink color) +(faint pink color) ++(obvious pink color) +++ (dark pink color)

#### Gene *27*; Score *0*

Images taken after 5 days at 37 ºC

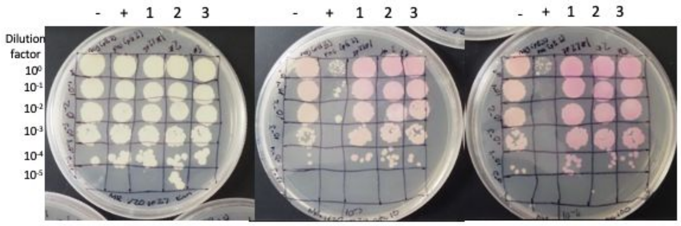


0 ng/ml aTc | 10 ng/ml aTc | 100 ng/ml aTc

| Lane | Gene ID | Plasmid name | Gene name | Toxic/Non-toxic | Colony color on 100 ng/ml aTc plate* |
|---|---|---|---|---|---|
| - Non-toxic control | -- | pExTra03 | Fruitloop *52* mutant | Non-toxic | + |
| + Toxic control | -- | pExTra02 | Fruitloop *52* | Toxic | - |
| 1 | | pExtra_27 | Amelie 27 replicate 1 | Non-toxic | ++ |
| 2 | | pExtra_27 | Amelie 27 replicate 2 | Non-toxic | ++ |
| 3 | | pExtra_27 | Amelie 27 replicate 3 | Non-toxic | ++ |

*Key: NG (no growth) - (no pink color) +(faint pink color) ++(obvious pink color) +++ (dark pink color)

#### Gene *28*; Score *0*

Images taken after 5 days at 37 ºC

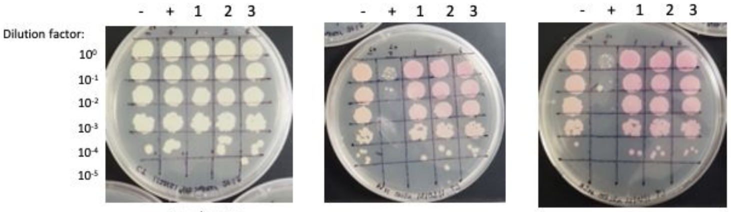


0 ng/ml aTc | 10 ng/ml aTc | 100 ng/ml aTc

| Lane | Gene ID | Plasmid name | Gene name | Toxic/Non-toxic | Colony color on 100 ng/ml aTc plate* |
|---|---|---|---|---|---|
| - Non-toxic control | -- | pExTra03 | Fruitloop *52* mutant | Non-toxic | ++ |
| + Toxic control | -- | pExTra02 | Fruitloop *52* | Toxic | - |
| 1 | -- | pExTra_Amelie28 | Amelie *28* replicate 1 | Non-toxic | ++ |
| 2 | -- | pExTra_Amelie28 | Amelie *28* replicate 2 | Non-toxic | ++ |
| 3 | -- | pExTra_Amelie28 | Amelie *28* replicate 3 | Non-toxic | ++ |

*Key: NG (no growth. )- (no pink color) +(faint pink color) ++(obvious pink color). +++ (dark pink color)

#### Gene *29*; Score *3*

Images taken after 5 days at 37 ºC

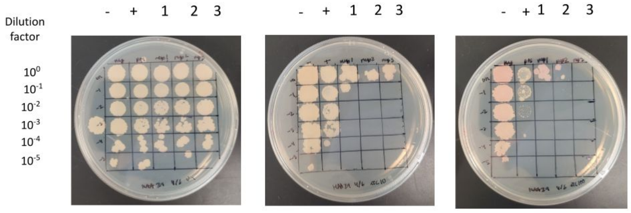


0 ng/ml aTc** | 10 ng/ml aTc | 100 ng/ml aTc

| Lane | Gene ID | Plasmid name | Gene name | Toxic/Non-toxic | Colony color on 100 ng/ml aTc plate* |
|---|---|---|---|---|---|
| - Non-toxic control | -- | pExTra03 | Fruitloop *52* mutant | Non-toxic | + |
| + Toxic control | -- | pExTra02 | Fruitloop *52* | Toxic | + |
| 1 | -- | pExTra_29 | Amelie *29* replicate 1 | Toxic | + |
| 2 | -- | pExTra_29 | Amelie *29* replicate 2 | Toxic | + |
| 3 | -- | pExTra_29 | Amelie *29* replicate 3 | Toxic | NG |

*Key: NG (no growth) - (no pink color) +(faint pink color) ++(obvious pink color) +++ (dark pink color)

#### Gene *30*; Score *0*

Images taken after 4 days at 37 ºC

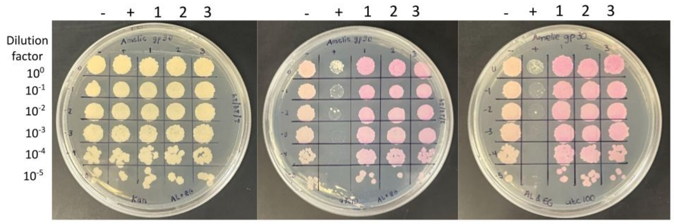


0 ng/ml aTc | 10 ng/ml aTc | 100 ng/ml aTc

| Lane | Gene ID | Plasmid name | Gene name | Toxic/Non-toxic | Colony color on 100 ng/ml aTc plate* |
|---|---|---|---|---|---|
| + Toxic control | -- | pExTra02 | Fruitloop *52* | Toxic | - |
| - Non-toxic control | -- | pExTra03 | Fruitloop *52* mutant | Non-toxic | + |
| 1 | 131436 | pExTra-Amelie30 | Amelie *30* replicate 1 | Non-toxic | ++ |
| 2 | 131436 | pExTra-Amelie30 | Amelie *30* replicate 2 | Non-toxic | ++ |
| 3 | 131436 | pExTra-Amelie30 | Amelie *30* replicate 3 | Non-toxic | ++ |

*Key: NG (no growth) - (no pink color) +(faint pink color) ++(obvious pink color) +++ (dark pink color)

#### Gene *31*; Score *2*

Images taken after 5 days at 37 ºC

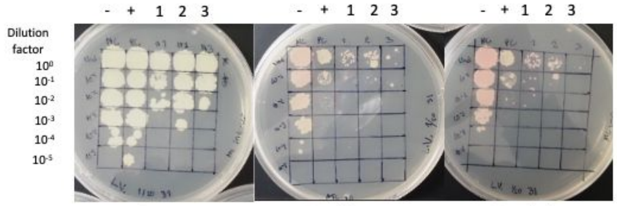


0 ng/ml aTc | 10 ng/ml aTc | 100 ng/ml aTc

| Lane | Gene ID | Plasmid name | Gene name | Toxic/Non-toxic | Colony color on 100 ng/ml aTc plate* |
|---|---|---|---|---|---|
| - Non-toxic control | -- | pExTra03 | Fruitloop *52* mutant | Non-toxic | ++ |
| + Toxic control | -- | pExTra02 | Fruitloop *52* | Toxic | - |
| 1 | -- | pExTra__Amelie31 | Amelie 31 replicate 1 | Toxic | + |
| 2 | -- | pExTra__Amelie31 | Amelie 31 replicate 2 | Toxic | + |
| 3 | -- | pExTra__Amelie31 | Amelie 31 replicate 3 | Toxic | NG |

*Key: NG (no growth) - (no pink color) +(faint pink color) ++(obvious pink color) +++ (dark pink color)

#### Gene *32*; Score *2*

Images taken after 4 days at 37 ºC

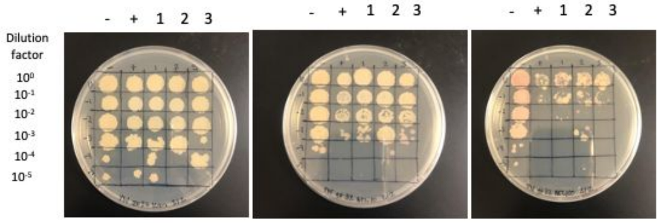


0 ng/ml aTc | 10 ng/ml aTc | 100 ng/ml aTc

| Lane | Gene ID | Plasmid name | Gene name | Toxic/Non-toxic | Colony color on 100 ng/ml aTc plate* |
|---|---|---|---|---|---|
| - Non-toxic control | -- | pExTra03 | Fruitloop *52* mutant | Non-toxic | ++ |
| + Toxic control | -- | pExTra02 | Fruitloop *52* | Toxic | - |
| 1 | -- | pExTra-Amelie 32 | Amelie 32 replicate 1 | toxic | + |
| 2 | -- | pExTra-Amelie 32 | Amelie 32 replicate 2 | toxic | + |
| 3 | -- | pExTra-Amelie 32 | Amelie 32 replicate 3 | toxic | + |

*Key: NG (no growth) - (no pink color) +(faint pink color)++(obvious pink color) +++ (dark pink color)

#### Gene *33*; Score *0*

Images taken after 5 days at 37 ºC

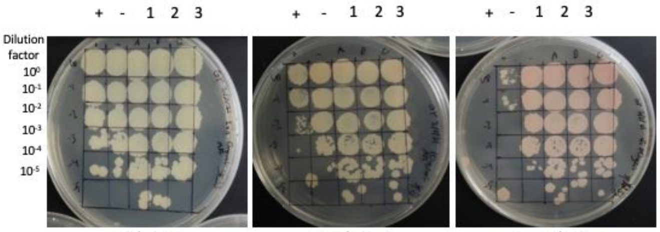


0 ng/ml aTc | 10 ng/ml aTc | 100 ng/ml aTc

| Lane | Gene ID | Plasmid name | Gene name | Toxic/Non-toxic | Colony color on 100 ng/ml aTc plate* |
|---|---|---|---|---|---|
| - Non-toxic control | -- | pExTra03 | Fruitloop *52* mutant | Non-toxic | ++ |
| + Toxic control | -- | pExTra02 | Fruitloop *52* | Toxic | + |
| 1 | -- | pExTra_33 | Amelie 33 replicate 1 | Non-toxic | ++ |
| 2 | -- | pExTra_33 | Amelie 33 replicate 2 | Non-toxic | ++ |
| 3 | -- | pExTra_33 | Amelie 33 replicate 3 | Non-toxic | ++ |

*Key: NG (no growth) - (no pink color) +(faint pink color) ++(obvious pink color) +++ (dark pink color)

#### Gene *34*; Score *3*

Images taken after 5 days at 37 ºC

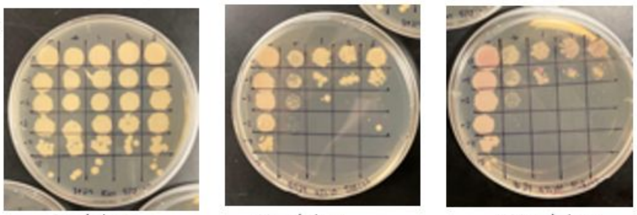


0 ng/ml aTc | 10 ng/ml aTc | 100 ng/ml aTc

| Lane | Gene ID | Plasmid name | Gene name | Toxic/Non-toxic | Colony color on 100 ng/ml aTc plate* |
|---|---|---|---|---|---|
| - Non-toxic control | -- | pExTra03 | Fruitloop *52* mutant | Non-toxic | + |
| + Toxic control | -- | pExTra02 | Fruitloop *52* | Toxic | - |
| 1 | | pExTra-Amelie 34 | Amelie 34 replicate 1 | Toxic | +++ |
| 2 | | pExTra-Amelie 34 | Amelie 34 replicate 2 | Toxic | ++ |
| 3 | | pExTra-Amelie 34 | Amelie 34 replicate 3 | Toxic | ++ |

*Key: NG (no growth) - (no pink color) +(faint pink color) ++(obvious pink color) +++ (dark pink color)

#### Gene *35*; Score *0*

Images taken after 5 days at 37 ºC

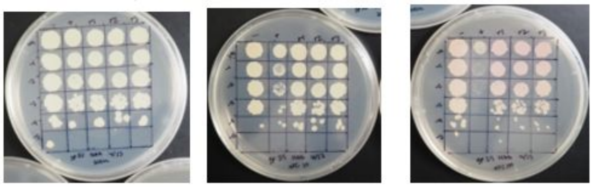


0 ng/ml aTc* | 10 ng/ml aTc | 100 ng/ml aTc

| Lane | Gene ID | Plasmid name | Gene name | Toxic/Non-toxic | Colony color on 100 ng/ml aTc plate* |
|---|---|---|---|---|---|
| - Non-toxic control | -- | pExTra03 | Fruitloop *52* mutant | Non-toxic | + |
| + Toxic control | -- | pExTra02 | Fruitloop *52* | Toxic | + |
| 1 | -- | pExTra_35 | Amelie *35* replicate 1 | Non-toxic | + |
| 2 | -- | pExTra_35 | Amelie *35* replicate 2 | Non-toxic | + |
| 3 | -- | pExTra_35 | Amelie *35* replicate 3 | Non-toxic | + |

*Key: NG (no growth) - (no pink color) +(faint pink color) ++(obvious pink color) +++ (dark pink color)

#### Gene *36*; Score *0*

Images taken after 5 days at 37 ºC

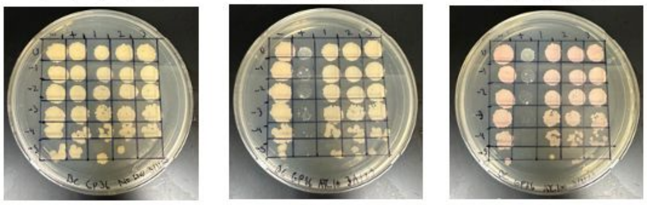


0 ng/ml aTc | 10 ng/ml aTc | 100 ng/ml aTc

| Lane | Gene ID | Plasmid name | Gene name | Toxic/Non-toxic | Colony color on 100 ng/ml aTc plate* |
|---|---|---|---|---|---|
| - Non-toxic control | -- | pExTra03 | Fruitloop *52* mutant | Non-toxic | + |
| + Toxic control | -- | pExTra02 | Fruitloop *52* | Toxic | - |
| 1 | -- | pExTra_36 | Amelie 36 replicate 1 | Non-toxic | + |
| 2 | -- | pExTra_36 | Amelie 36 replicate 2 | Non-toxic | + |
| 3 | -- | pExTra_36 | Amelie 36 replicate 3 | Non-toxic | ++ |

*Key: NG (no growth) - (no pink color) +(faint pink color) ++(obvious pink color) +++ (dark pink color)

#### Gene *37*; Score *0*

Images taken after 4 days at 37 ºC

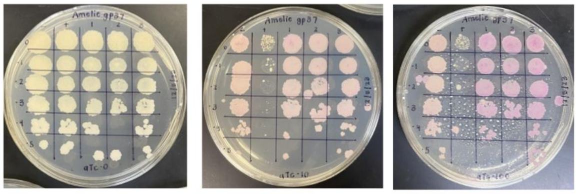


0 ng/ml aTc | 10 ng/ml aTc | 100 ng/ml aTc

| Lane | Gene ID | Plasmid name | Gene name | Toxic/Non-toxic | Colony color on 100 ng/ml aTc plate* |
|---|---|---|---|---|---|
| - Non-toxic control | -- | pExTra03 | Fruitloop *52* mutant | Non-toxic | ++ |
| + Toxic control | -- | pExTra02 | Fruitloop *52* | Toxic | - |
| 1 | | pExTra-Amelie37 | Amelie *37*replicate 1 | Non-toxic | +++ |
| 2 | | pExTra-Amelie37 | Amelie *37* replicate 2 | Non-toxic | +++ |
| 3 | | pExTra-Amelie37 | Amelie *37* replicate 3 | Non-toxic | +++ |

*Key: NG (no growth) - (no pink color) +(faint pink color) ++(obvious pink color) +++ (dark pink color)

#### Gene *38*; Score *0*

Images taken after 4 days at 37 ºC

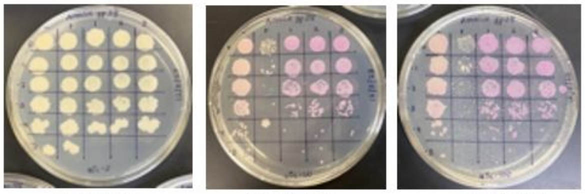


0 ng/ml aTc | 10 ng/ml aTc | 100 ng/ml aTc

| Lane | Gene ID | Plasmid name | Gene name | Toxic/Non-toxic | Colony color on 100 ng/ml aTc plate* |
|---|---|---|---|---|---|
| - Non-toxic control | -- | pExTra03 | Fruitloop *52* mutant | Non-toxic | ++ |
| + Toxic control | -- | pExTra02 | Fruitloop *52* | Toxic | - |
| 1 | | pExTra-Amelie38 | Amelie *38* replicate 1 | Non-toxic | +++ |
| 2 | | pExTra-Amelie38 | Amelie *38* replicate 2 | Non-toxic | +++ |
| 3 | | pExTra-Amelie38 | Amelie *38* replicate 3 | Non-toxic | +++ |

*Key: NG (no growth) - (no pink color) +(faint pink color) ++(obvious pink color) +++ (dark pink color)

#### Gene *39*; Score *0*

Images taken after 5 days at 37 ºC

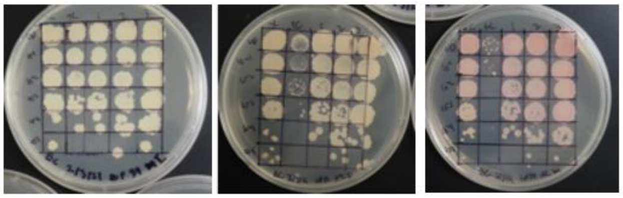


0 ng/ml aTc | 10 ng/ml aTc | 100 ng/ml aTc

| Lane | Gene ID | Plasmid name | Gene name | Toxic/Non-toxic | Colony color on 100 ng/ml aTc plate* |
|---|---|---|---|---|---|
| - Non-toxic control | -- | pExTra03 | Fruitloop *52* mutant | Non-toxic | + |
| + Toxic control | -- | pExTra02 | Fruitloop *52* | Toxic | - |
| 1 | -- | pExTra-39 | Amelie 39 replicate 1 | Non-toxic | ++ |
| 2 | -- | pExTra-39 | Amelie 39 replicate 2 | Non-toxic | ++ |
| 3 | -- | pExTra-39 | Amelie 39 replicate 3 | Non-toxic | ++ |

*Key: NG (no growth) - (no pink color) +(faint pink color) ++(obvious pink color) +++ (dark pink color)

#### Gene *40*; Score *0*

Images taken after 5 days at 37 ºC

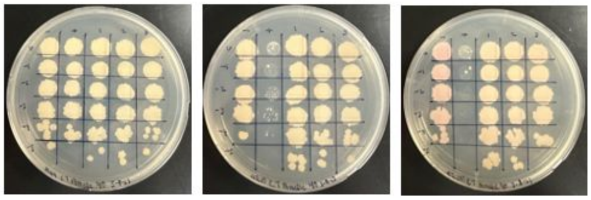


0 ng/ml aTc | 10 ng/ml aTc | 100 ng/ml aTc

| Lane | Gene ID | Plasmid name | Gene name | Toxic/Non-toxic | Colony color on 100 ng/ml aTc plate* |
|---|---|---|---|---|---|
| - Non-toxic control | -- | pExTra03 | Fruitloop *52* mutant | Non-toxic | + |
| + Toxic control | -- | pExTra02 | Fruitloop *52* | Toxic | - |
| 1 | | pExtra_40 | Amelie 40 replicate 1 | Non-toxic | + |
| 2 | | pExtra_40 | Amelie 40 replicate 2 | Non-toxic | + |
| 3 | | pExtra_40 | Amelie 40 replicate 3 | Non-toxic | + |

*Key: NG (no growth) - (no pink color) +(faint pink color) ++(obvious pink color) +++ (dark pink color)

#### Gene *41*; Score *0*

Images taken after 5 days at 37ºC

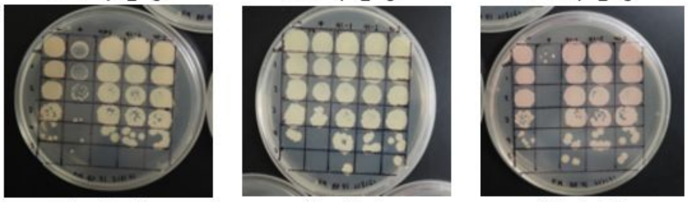


0 ng/ml aTc | 10 ng/ml aTc | 100 ng/ml aTc

| Lane | Gene ID | Plasmid Name | Gene Name | Toxic/Non-Toxic | Colony Color on 100 ng/mL aTc Plate |
|---|---|---|---|---|---|
| (-) Non-Toxic Control | ----- | pExTra_03 | Fruitloop *52* Mutant | Non-Toxic | + |
| (+) Toxic Control | ----- | pExTra_02 | Fruitloop *52* | Toxic | - |
| 1 | ----- | pExTra_41 | Amelie 41 Replicate 1 | Non-Toxic | + |
| 2 | ----- | pExTra_41 | Amelie 41 Replicate 2 | Non-Toxic | + |
| 3 | ----- | pExTra_41 | Amelie 41 Replicate 3 | Non-Toxic | + |

*Key: NG (No Growth) - (no pink color) + (faint pink color) ++ (obvious pink color) +++ (dark pink color)

Images taken after 5 days at 37 ºC

#### Gene *42*; Score *0*

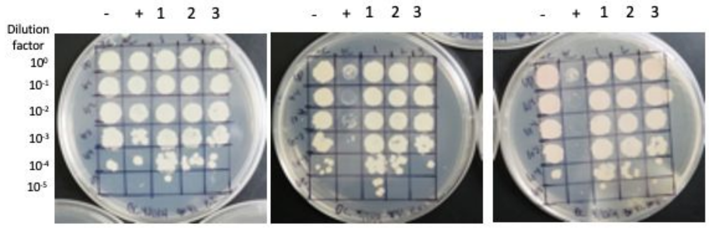


0 ng/ml aTc | 10 ng/ml aTc | 100 ng/ml aTc

| Lane | Gene ID | Plasmid name | Gene name | Toxic/Non-toxic | Colony color on 100 ng/ml aTc plate* |
|---|---|---|---|---|---|
| - Non-toxic control | -- | pExTra03 | Fruitloop *52* mutant | Non-toxic | + |
| + Toxic control | -- | pExTra02 | Fruitloop *52* | Toxic | - |
| 1 | -- | pExTra_42 | Amelie *42* replicate 1 | Non-toxic | + |
| 2 | -- | pExTra_42 | Amelie *42* replicate 2 | Non-toxic | + |
| 3 | -- | pExTra_42 | Amelie *42* replicate 3 | Non-toxic | + |

*Key: NG (no growth) - (no pink color) +(faint pink color) ++(obvious pink color) +++ (dark pink color)

Images taken after 5 days at 37 ºC

#### Gene *43*; Score *3*

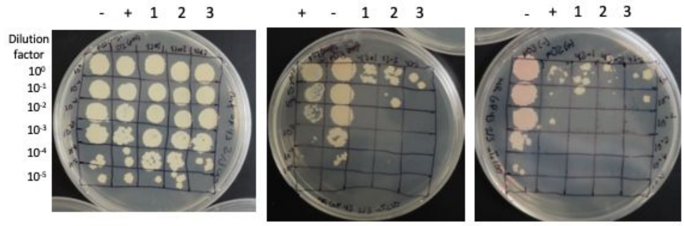


0 ng/ml aTc | 10 ng/ml aTc | 100 ng/ml aTc

| Lane | Gene ID | Plasmid name | Gene name | Toxic/Non-toxic | Colony color on 100 ng/ml aTc plate* |
|---|---|---|---|---|---|
| - Non-toxic control | -- | pExTra03 | Fruitloop *52* mutant | Non-toxic | + |
| + Toxic control | -- | pExTra02 | Fruitloop *52* | Toxic | - |
| 1 | | pExtra_43 | Amelie 43 replicate 1 | Toxic | - |
| 2 | | pExtra_43 | Amelie 43 replicate 2 | Toxic | - |
| 3 | | pExtra_43 | Amelie 43 replicate 3 | Toxic | - |

*Key: NG (no growth) - (no pink color) +(faint pink color) ++(obvious pink color) +++ (dark pink color)

Images taken after 4 days at 37 ºC

#### Gene *44*; Score *3*

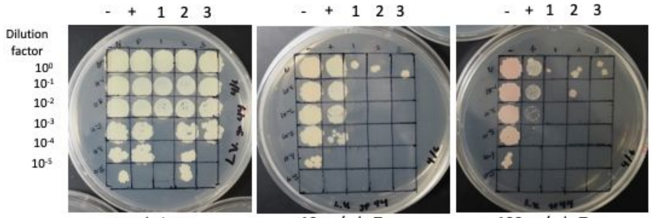


0 ng/ml aTc | 10 ng/ml aTc | 100 ng/ml aTc

| Lane | Gene ID | Plasmid name | Gene name | Toxic/Non-toxic | Colony color on 100 ng/ml aTc plate* |
|---|---|---|---|---|---|
| - Non-toxic control | -- | pExTra03 | Fruitloop *52* mutant | Non-toxic | ++ |
| + Toxic control | -- | pExTra02 | Fruitloop *52* | Toxic | - |
| 1 | -- | pExTra__44 | Amelie 44 replicate 1 | Toxic | NG |
| 2 | -- | pExTra__44 | Amelie 44 replicate 1 | Toxic | + |
| 3 | -- | pExTra__44 | Amelie 44 replicate 1 | Toxic | + |

*Key: NG (no growth) - (no pink color) +(faint pink color) ++(obvious pink color) +++

Images taken after 5 days at 37 ºC

#### Gene *45*; Score *0*

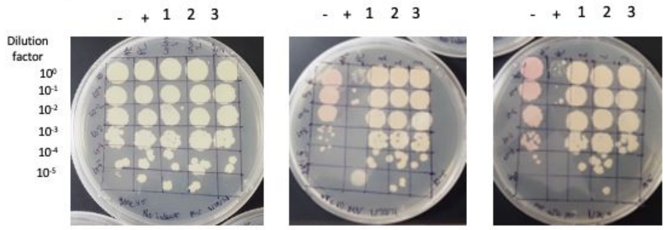


0 ng/ml aTc | 10 ng/ml aTc | 100 ng/ml aTc

| Lane | Gene ID | Plasmid name | Gene name | Toxic/Non-toxic | Colony color on 100 ng/ml aTc plate* |
|---|---|---|---|---|---|
| - Non-toxic control | -- | pExTra03 | Fruitloop *52* mutant | Non-toxic | ++ |
| + Toxic control | -- | pExTra02 | Fruitloop *52* | Toxic | - |
| 1 | -- | pExTra_45 | Amelie *45* replicate 1 | Non-toxic | + |
| 2 | -- | pExTra_45 | Amelie *45* replicate 2 | Non-toxic | + |
| 3 | -- | pExTra_45 | Amelie *45* replicate 3 | Non-toxic | + |

*Key: NG (no growth) - (no pink color) +(faint pink color) ++(obvious pink color) +++ (dark pink color)

Images taken after 5 days at 37 ºC

#### Gene *46*; Score *1*

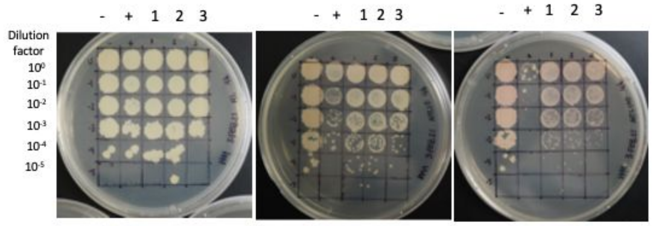


0 ng/ml aTc | 10 ng/ml aTc | 100 ng/ml aTc

| Lane | Gene ID | Plasmid name | Gene name | Toxic/Non-toxic | Colony color on 100 ng/ml aTc plate* |
|---|---|---|---|---|---|
| - Non-toxic control | -- | pExTra03 | Fruitloop *52* mutant | Non-toxic | ++ |
| + Toxic control | -- | pExTra02 | Fruitloop *52* | Toxic | + |
| 1 | -- | pExTra_46 | Amelie *46* replicate 1 | toxic | + |
| 2 | -- | pExTra_46 | Amelie *46* replicate 2 | toxic | + |
| 3 | -- | pExTra_46 | Amelie *46* replicate 3 | toxic | + |

*Key: NG (no growth) - (no pink color) +(faint pink color) ++(obvious pink color) +++ (dark pink color)

Images taken after 5 days at 37 ºC

#### Gene *47*; Score *3*

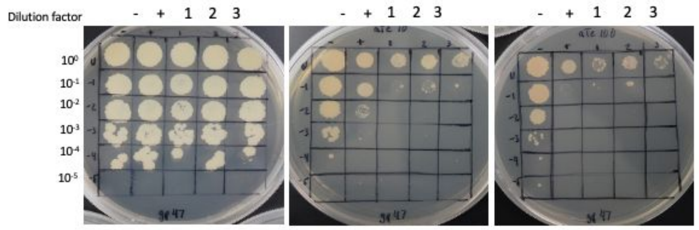


0 ng/ml aTc | 10 ng/ml aTc | 100 ng/ml aTc

| Lane | Gene ID | Plasmid name | Gene name | Toxic/Non-toxic | Colony color on 100 ng/ml aTc plate* |
|---|---|---|---|---|---|
| - Non-toxic control | -- | pExTra03 | Fruitloop *52* mutant | Non-toxic | + |
| + Toxic control | -- | pExTra02 | Fruitloop *52* | Toxic | - |
| 1 | | pExTra- Amelie 47 | Amelie 47 replicate 1 | Toxic | - |
| 2 | | pExTra- Amelie 47 | Amelie 47 replicate 2 | Toxic | - |
| 3 | | pExTra- Amelie 47 | Amelie 47replicate 3 | Toxic | - |

*Key: NG (no growth) - (no pink color) +(faint pink color) ++(obvious pink color) +++ (dark pink color)

Images taken after 4 days at 37 ºC

#### Gene *48*; Score *0*

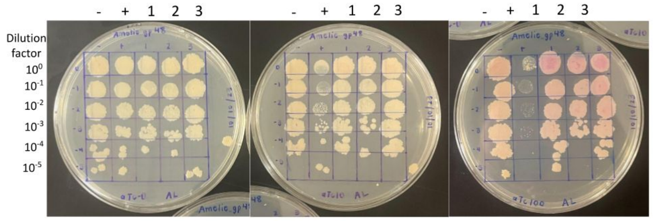


0 ng/ml aTc | 10 ng/ml aTc | 100 ng/ml aTc

| Lane | Gene ID | Plasmid name | Gene name | Toxic/Non-toxic | Colony color on 100 ng/ml aTc plate* |
|---|---|---|---|---|---|
| + Toxic control | -- | pExTra02 | Fruitloop *52* | Toxic | - |
| - Non-toxic control | -- | pExTra03 | Fruitloop *52* mutant | Non-toxic | + |
| 1 | 131436 | pExTra-Amelie48 | Amelie *48* replicate 1 | Non-toxic | ++ |
| 2 | 131436 | pExTra-Amelie48 | Amelie *48* replicate 2 | Non-toxic | ++ |
| 3 | 131436 | pExTra-Amelie48 | Amelie *48* replicate 3 | Non-toxic | ++ |

*Key: NG (no growth) - (no pink color) +(faint pink color) ++(obvious pink color) +++ (dark pink color)

Images taken after 4 days at 37 ºC

#### Gene *49*; Score *3*

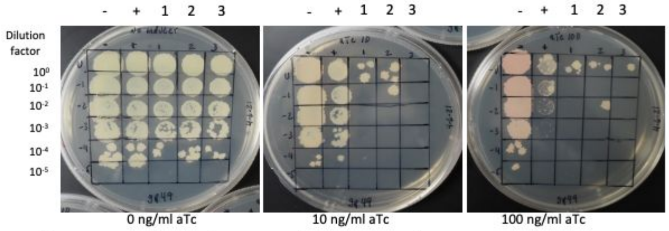


0 ng/ml aTc | 10 ng/ml aTc | 100 ng/ml aTc

| Lane | Gene ID | Plasmid name | Gene name | Toxic/Non-toxic | Colony color on 100 ng/ml aTc plate* |
|---|---|---|---|---|---|
| - Non-toxic control | -- | pExTra03 | Fruitloop *52* mutant | Non-toxic | + |
| + Toxic control | -- | pExTra02 | Fruitloop *52* | Toxic | - |
| 1 | -- | pExTra-Amelie 49 | Amelie 49 replicate 1 | Toxic | - |
| 2 | -- | pExTra-Amelie 49 | Amelie 49 replicate 2 | Toxic | - |
| 3 | -- | pExTra-Amelie 49 | Amelie 49 replicate 3 | Toxic | - |

*Key: NG (no growth) - (no pink color) +(faint pink color)++(obvious pink color) +++ (dark pink color)

#### Gene *50*; Score 2

Images taken after 5 days at 37 ºC

- + 1 2 3    - + 1 2 3    - + 1 2 3

Dilution factor: $10^{0}$, $10^{-1}$, $10^{-2}$, $10^{-3}$, $10^{-4}$, $10^{-5}$

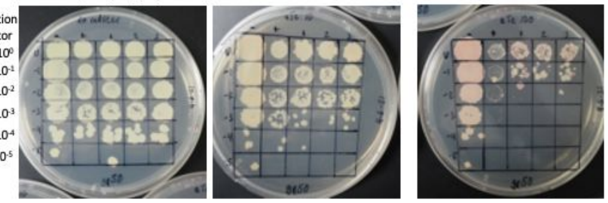

0 ng/ml aTc    10 ng/ml aTc    100 ng/ml aTc

| Lane | Gene ID | Plasmid name | Gene name | Toxic/Non-toxic | Colony color on 100 ng/ml aTc plate* |
|---|---|---|---|---|---|
| - Non-toxic control | -- | pExTra03 | Fruitloop *52* mutant | Non-toxic | ++ |
| + Toxic control | -- | pExTra02 | Fruitloop *52* | Toxic | + |
| 1 | -- | pExTra_50 | Amelie *50* replicate 1 | Toxic | + |
| 2 | -- | pExTra_50 | Amelie *50* replicate 2 | Toxic | + |
| 3 | -- | pExTra_50 | Amelie *50* replicate 3 | Toxic | + |

*Key: NG (no growth) - (no pink color) +(faint pink color) ++(obvious pink color) +++ (dark pink color)

#### Gene *51*; Score 0

Images taken after 5 days at 37 ºC

Dilution factor   - + 1 2 3    - + 1 2 3    - + 1 2 3

$10^{0}$, $10^{-1}$, $10^{-2}$, $10^{-3}$, $10^{-4}$, $10^{-5}$

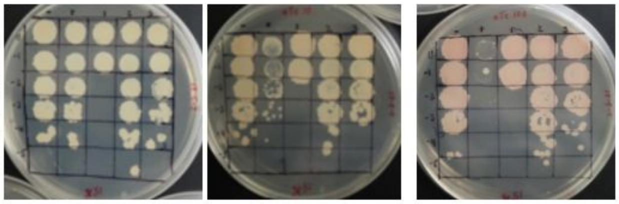

0 ng/ml aTc    10 ng/ml aTc    100 ng/ml aTc

| Lane | Gene ID | Plasmid name | Gene name | Toxic/Non-toxic | Colony color on 100 ng/ml aTc plate* |
|---|---|---|---|---|---|
| - Non-toxic control | -- | pExTra03 | Fruitloop *52* mutant | Non-toxic | ++ |
| + Toxic control | -- | pExTra02 | Fruitloop *52* | Toxic | - |
| 1 | | pExTra- Amelie 51 | Amelie 51 replicate 1 | Non-toxic | ++ |
| 2 | | pExTra- Amelie 51 | Amelie 51 replicate 2 | Non-toxic | ++ |
| 3 | | pExTra- Amelie 51 | Amelie 51 replicate 3 | Non-toxic | ++ |

*Key: NG (no growth) - (no pink color) +(faint pink color) ++(obvious pink color) +++ (dark pink color)

#### Gene *52*; Score 0

Images taken after 5 days at 37 ºC

- + 1 2 3    - + 1 2 3    - + 1 2 3

Dilution factor: $10^{0}$, $10^{-1}$, $10^{-2}$, $10^{-3}$, $10^{-4}$, $10^{-5}$

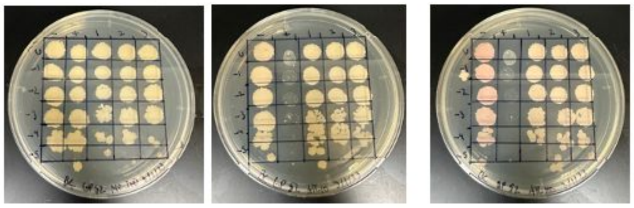

0 ng/ml aTc    10 ng/ml aTc    100 ng/ml aTc

| Lane | Gene ID | Plasmid name | Gene name | Toxic/Non-toxic | Colony color on 100 ng/ml aTc plate* |
|---|---|---|---|---|---|
| - Non-toxic control | -- | pExTra03 | Fruitloop *52* mutant | Non-toxic | + |
| + Toxic control | -- | pExTra02 | Fruitloop *52* | Toxic | - |
| 1 | -- | pExTra_52 | Amelie *52* replicate 1 | Non-toxic | + |
| 2 | -- | pExTra_52 | Amelie *52* replicate 2 | Non-toxic | + |
| 3 | -- | pExTra_52 | Amelie *52* replicate 3 | Non-toxic | + |

*Key: NG (no growth) - (no pink color) +(faint pink color) ++(obvious pink color) +++ (dark pink color)

#### Gene *53*; Score 0

Images taken after 4 days at 37 ºC

- + 1 2 3    - + 1 2 3    - + 1 2 3

Dilution factor: $10^{0}$, $10^{-1}$, $10^{-2}$, $10^{-3}$, $10^{-4}$, $10^{-5}$

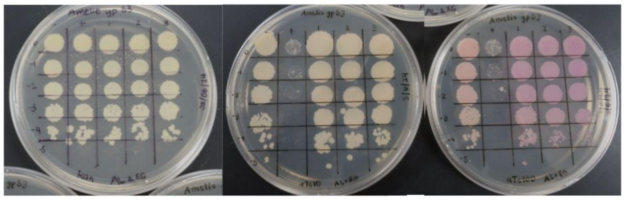

0 ng/ml aTc    10 ng/ml aTc    100 ng/ml aTc

| Lane | Gene ID | Plasmid name | Gene name | Toxic/Non-toxic | Colony color on 100 ng/ml aTc plate* |
|---|---|---|---|---|---|
| + Toxic control | -- | pExTra02 | Fruitloop *52* | Toxic | - |
| - Non-toxic control | -- | pExTra03 | Fruitloop *52* mutant | Non-toxic | + |
| 1 | | pExTra-Amelie53 | Amelie *53* replicate 1 | Non-toxic | ++ |
| 2 | | pExTra-Amelie53 | Amelie *53* replicate 2 | Non-toxic | ++ |
| 3 | | pExTra-Amelie53 | Amelie *53* replicate 3 | Non-toxic | ++ |

*Key: NG (no growth) - (no pink color) +(faint pink color) ++(obvious pink color) +++ (dark pink color)

#### Gene *54*; Score 0

Images taken after 5 days at 37 ºC

- + 1 2 3    - + 1 2 3    - + 1 2 3

Dilution factor: 10, 10-, 10-, 10-, 10-, 10-

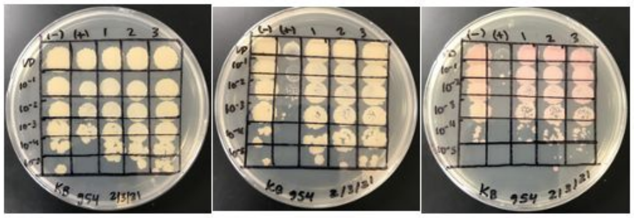

0 ng/ml aTc    10 ng/ml aTc    100 ng/ml aTc

| Lane | Gene ID | Plasmid name | Gene name | Toxic/Non-toxic | Colony color on 100 ng/ml aTc plate* |
|---|---|---|---|---|---|
| - Non-toxic control | -- | pExTra03 | Fruitloop *52* mutant | Non-toxic | ++ |
| + Toxic control | -- | pExTra02 | Fruitloop *52* | Toxic | - |
| 1 | -- | pExTra_54 | Amelie 54 replicate 1 | Non-toxic | + |
| 2 | -- | pExTra_54 | Amelie 54 replicate 2 | Non-toxic | + |
| 3 | -- | pExTra_54 | Amelie 54 replicate 3 | Non-toxic | + |

*Key: NG (no growth) - (no pink color) +(faint pink color) ++(obvious pink color) +++

#### Gene *55*; Score 1

Images taken after 4 days at 37 ºC

- + 1 2 3    - + 1 2 3    - + 1 2 3

Dilution factor: $10^{0}$, $10^{-1}$, $10^{-2}$, $10^{-3}$, $10^{-4}$, $10^{-5}$

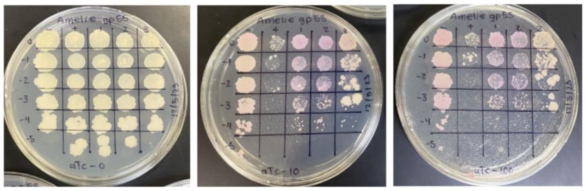

0 ng/ml aTc    10 ng/ml aTc    100 ng/ml aTc

| Lane | Gene ID | Plasmid name | Gene name | Toxic/Non-toxic | Colony color on 100 ng/ml aTc plate* |
|---|---|---|---|---|---|
| - Non-toxic control | -- | pExTra03 | Fruitloop *52* mutant | Non-toxic | ++ |
| + Toxic control | -- | pExTra02 | Fruitloop *52* | Toxic | - |
| 1 | | pExTra-Amelie55 | Amelie *55* replicate 1 | Toxic | +++ |
| 2 | | pExTra-Amelie55 | Amelie *55* replicate 2 | Toxic | +++ |
| 3 | | pExTra-Amelie55 | Amelie *55* replicate 3 | Toxic | +++ |

*Key: NG (no growth) - (no pink color) +(faint pink color) ++(obvious pink color) +++ (dark pink color)

#### Gene *56*; Score 1

Images taken after 5 days at 37 ºC

- + 1 2 3    - + 1 2 3    - + 1 2 3

Dilution factor: $10^{0}$, $10^{-1}$, $10^{-2}$, $10^{-3}$, $10^{-4}$, $10^{-5}$

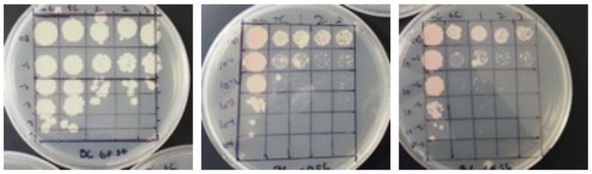

0 ng/ml aTc    10 ng/ml aTc    100 ng/ml aTc

| Lane | Gene ID | Plasmid name | Gene name | Toxic/Non-toxic | Colony color on 100 ng/ml aTc plate* |
|---|---|---|---|---|---|
| - Non-toxic control | -- | pExTra03 | Fruitloop *52* mutant | Non-toxic | ++ |
| + Toxic control | -- | pExTra02 | Fruitloop *52* | Toxic | - |
| 1 | -- | pExTra_56 | Amelie *56* replicate 1 | Toxic | + |
| 2 | -- | pExTra_56 | Amelie *56* replicate 2 | Toxic | + |
| 3 | -- | pExTra_56 | Amelie *56* replicate 3 | Toxic | + |

*Key: NG (no growth) - (no pink color) +(faint pink color) ++(obvious pink color) +++ (dark pink color)

#### Gene *57*; Score 3

Images taken after 5 days at 37 ºC

- + 1 2 3    - + 1 2 3    - + 1 2 3

Dilution factor: $10^{0}$, $10^{-1}$, $10^{-2}$, $10^{-3}$, $10^{-4}$, $10^{-5}$

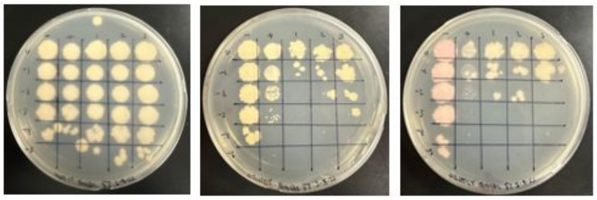

0 ng/ml aTc    10 ng/ml aTc    100 ng/ml aTc

| Lane | Gene ID | Plasmid name | Gene name | Toxic/Non-toxic | Colony color on 100 ng/ml aTc plate* |
|---|---|---|---|---|---|
| - Non-toxic control | -- | pExTra03 | Fruitloop *52* mutant | Non-toxic | + |
| + Toxic control | -- | pExTra02 | Fruitloop *52* | Toxic | - |
| 1 | | pExtra_57 | Amelie 57 replicate 1 | Toxic | + |
| 2 | | pExtra_57 | Amelie 57 replicate 2 | Toxic | + |
| 3 | | pExtra_57 | Amelie 57 replicate 3 | Toxic | + |

*Key: NG (no growth) - (no pink color) +(faint pink color) ++(obvious pink color) +++ (dark pink color)

### Gene *58*; Score *1*

Images taken after 5 days at 37 ºC

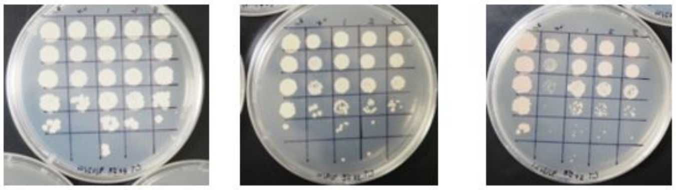


0 ng/ml aTc 10 ng/ml aTc 100 ng/ml aTc

| Lane | Gene ID | Plasmid name | Gene name | Toxic/Non-toxic | Colony color on 100 ng/ml aTc plate* |
|---|---|---|---|---|---|
| - Non-toxic control | -- | pExTra03 | Fruitloop *52* mutant | Non-toxic | + |
| + Toxic control | -- | pExTra02 | Fruitloop *52* | Toxic | - |
| 1 | -- | pExTra_58 | Amelie *58* replicate 1 | Toxic | + |
| 2 | -- | pExTra_58 | Amelie *58* replicate 2 | Toxic | + |
| 3 | -- | pExTra_58 | Amelie *58* replicate 3 | Toxic | + |

*Key: NG (no growth) - (no pink color) +(faint pink color) ++(obvious pink color) +++ (dark pink color)

### Gene *59*; Score *0*

Images taken after 5 days at 37°C

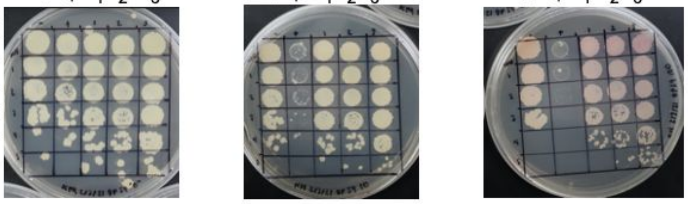


0 ng/ml aTc 10 ng/ml aTc 100 ng/ml aTc

| Lane | Gene ID | Plasmid Name | Gene Name | Toxic/Non-Toxic | Colony Color on 100 ng/mL aTc Plate |
|---|---|---|---|---|---|
| (-) Non-Toxic Control | ----- | pExTra_03 | Fruitloop *52* Mutant | Non-Toxic | + |
| (+) Toxic Control | ----- | pExTra_02 | Fruitloop *52* | Toxic | - |
| 1 | ----- | pExTra_59 | Amelie 59 Replicate 1 | Non-Toxic | + |
| 2 | ----- | pExTra_59 | Amelie 59 Replicate 2 | Non-Toxic | + |
| 3 | ----- | pExTra_59 | Amelie 59 Replicate 3 | Non-Toxic | ++ |

*Key: NG (No Growth) - (no pink color) + (faint pink color) ++ (obvious pink color) +++ (dark pink color)

### Gene *60*; Score *0*

Images taken after 5 days at 37 ºC

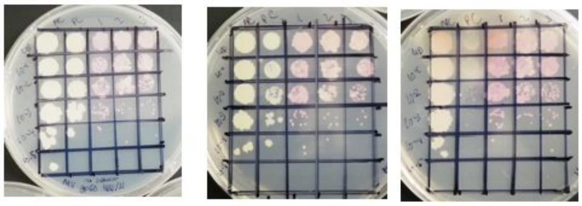


0 ng/ml aTc 10 ng/ml aTc 100 ng/ml aTc

| Lane | Gene ID | Plasmid name | Gene name | Toxic/Non-toxic | Colony color on 100 ng/ml aTc plate* |
|---|---|---|---|---|---|
| - Non-toxic control | -- | pExTra03 | Fruitloop *52* mutant | Non-toxic | + |
| + Toxic control | -- | pExTra02 | Fruitloop *52* | Toxic | - |
| 1 | -- | pExTra_60 | Amelie *60* replicate 1 | Non-toxic | ++ |
| 2 | -- | pExTra_60 | Amelie *60* replicate 2 | Non-toxic | ++ |
| 3 | -- | pExTra_60 | Amelie *60* replicate 3 | Non-toxic | ++ |

*Key: NG (no growth) - (no pink color) +(faint pink color) ++(obvious pink color) +++ (dark pink color)

### Gene *61*; Score *0*

Images taken after 5 days at 37 ºC

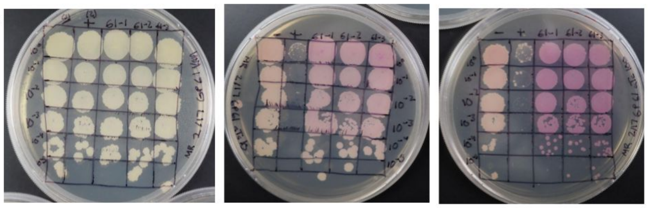


0 ng/ml aTc 10 ng/ml aTc 100 ng/ml aTc

| Lane | Gene ID | Plasmid name | Gene name | Toxic/Non-toxic | Colony color on 100 ng/ml aTc plate* |
|---|---|---|---|---|---|
| - Non-toxic control | -- | pExTra03 | Fruitloop *52* mutant | Non-toxic | + |
| + Toxic control | -- | pExTra02 | Fruitloop *52* | Toxic | - |
| 1 | | pExtra-Amelie_61 | Amelie 61 replicate 1 | Non-toxic | +++ |
| 2 | | pExtra-Amelie_61 | Amelie 61 replicate 2 | Non-toxic | +++ |
| 3 | | pExtra-Amelie_61 | Amelie 61 replicate 3 | Non-toxic | +++ |

*Key: NG (no growth) - (no pink color) +(faint pink color) ++(obvious pink color) +++ (dark pink color)

### Gene *62*; Score *2*

Images taken after 5 days at 37 ºC

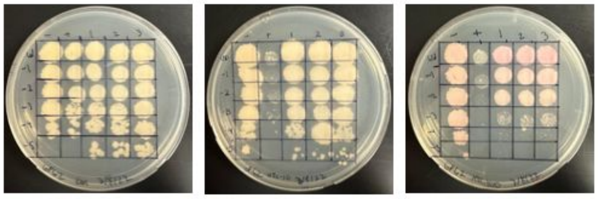


0 ng/ml aTc 10 ng/ml aTc 100 ng/ml aTc

| Lane | Gene ID | Plasmid name | Gene name | Toxic/Non-toxic | Colony color on 100 ng/ml aTc plate* |
|---|---|---|---|---|---|
| - Non-toxic control | -- | pExTra03 | Fruitloop *52* mutant | Non-toxic | + |
| + Toxic control | -- | pExTra02 | Fruitloop *52* | Toxic | - |
| 1 | | pExtra_62 | Amelie 62 replicate 1 | Toxic | ++ |
| 2 | | pExtra_62 | Amelie 62 replicate 2 | Toxic | ++ |
| 3 | | pExtra_62 | Amelie 62 replicate 3 | Toxic | ++ |

*Key: NG (no growth) - (no pink color) +(faint pink color) ++(obvious pink color) +++ (dark pink color)

### Gene *63*; Score *1*

Images taken after 4 days at 37 ºC

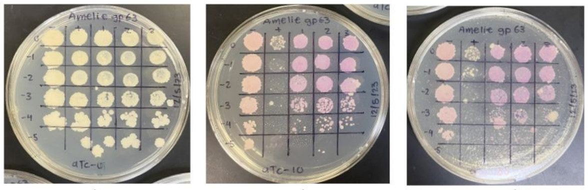


0 ng/ml aTc 10 ng/ml aTc 100 ng/ml aTc

| Lane | Gene ID | Plasmid name | Gene name | Toxic/Non-toxic | Colony color on 100 ng/ml aTc plate* |
|---|---|---|---|---|---|
| - Non-toxic control | -- | pExTra03 | Fruitloop *52* mutant | Non-toxic | ++ |
| + Toxic control | -- | pExTra02 | Fruitloop *52* | Toxic | - |
| 1 | | pExTra-Amelie63 | Amelie *63* replicate 1 | Toxic | +++ |
| 2 | | pExTra-Amelie63 | Amelie *63* replicate 2 | Toxic | +++ |
| 3 | | pExTra-Amelie63 | Amelie *63* replicate 3 | Toxic | +++ |

*Key: NG (no growth) - (no pink color) +(faint pink color) ++(obvious pink color) +++ (dark pink color)

### Gene *64*; Score *0*

Images taken after 5 days at 37 ºC

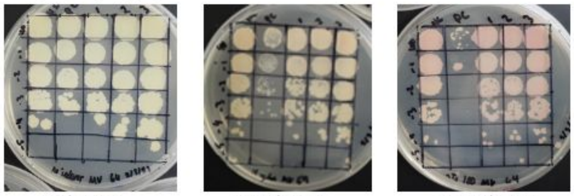


0 ng/ml aTc 10 ng/ml aTc 100 ng/ml aTc

| Lane | Gene ID | Plasmid name | Gene name | Toxic/Non-toxic | Colony color on 100 ng/ml aTc plate* |
|---|---|---|---|---|---|
| - Non-toxic control | -- | pExTra03 | Fruitloop *52* mutant | Non-toxic | ++ |
| + Toxic control | -- | pExTra02 | Fruitloop *52* | Toxic | - |
| 1 | -- | pExTra_64 | Amelie *64* replicate 1 | Non-toxic | ++ |
| 2 | -- | pExTra_64 | Amelie *64* replicate 2 | Non-toxic | ++ |
| 3 | -- | pExTra_64 | Amelie *64* replicate 3 | Non-toxic | ++ |

*Key: NG (no growth) - (no pink color) +(faint pink color) ++(obvious pink color) +++ (dark pink color)

### Gene *65*; Score *1*

Images taken after 4 days at 37 ºC

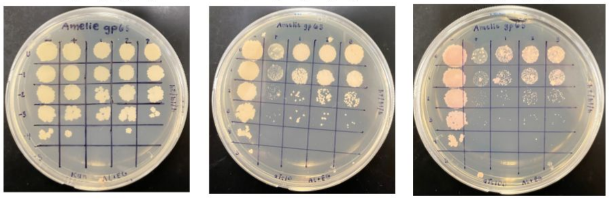


0 ng/ml aTc 10 ng/ml aTc 100 ng/ml aTc

| Lane | Gene ID | Plasmid name | Gene name | Toxic/Non-toxic | Colony color on 100 ng/ml aTc plate* |
|---|---|---|---|---|---|
| + Toxic control | -- | pExTra02 | Fruitloop *52* | Toxic | + |
| - Non-toxic control | -- | pExTra03 | Fruitloop *52* mutant | Non-toxic | - |
| 1 | 131436 | pExTra-Amelie65 | Amelie *65* replicate 1 | Toxic | + |
| 2 | 131436 | pExTra-Amelie65 | Amelie *65* replicate 2 | Toxic | + |
| 3 | 131436 | pExTra-Amelie65 | Amelie *65* replicate 3 | Toxic | + |

*Key: NG (no growth) - (no pink color) +(faint pink color) ++(obvious pink color) +++ (dark pink color)

Images taken after 5 days at 37 ºC

#### Gene *66*; Score *0*

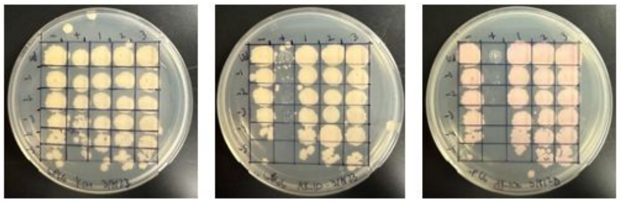


0 ng/ml aTc | 10 ng/ml aTc | 100 ng/ml aTc

| Lane | Gene ID | Plasmid name | Gene name | Toxic/Non-toxic | Colony color on 100 ng/ml aTc plate* |
|---|---|---|---|---|---|
| - Non-toxic control | -- | pExTra03 | Fruitloop *52* mutant | Non-toxic | + |
| + Toxic control | -- | pExTra02 | Fruitloop *52* | Toxic | - |
| 1 | | pExtra_66 | Amelie 66 replicate 1 | Non-toxic | ++ |
| 2 | | pExtra_66 | Amelie 66 replicate 2 | Non-toxic | ++ |
| 3 | | pExtra_66 | Amelie 66 replicate 3 | Non-toxic | ++ |

*Key: NG (no growth) - (no pink color) +(faint pink color) ++(obvious pink color) +++ (dark pink color)

Images taken after 5 days at 37 ºC

#### Gene *67*; Score *0*

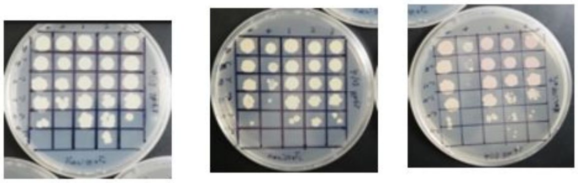


0 ng/ml aTc | 10 ng/ml aTc | 100 ng/ml aTc

| Lane | Gene ID | Plasmid name | Gene name | Toxic/Non-toxic | Colony color on 100 ng/ml aTc plate* |
|---|---|---|---|---|---|
| - Non-toxic control | -- | pExTra03 | Fruitloop *52* mutant | Non-toxic | + |
| + Toxic control | -- | pExTra02 | Fruitloop *52* | Toxic | - |
| 1 | -- | pExTra_67 | Amelie *67* replicate 1 | Non-toxic | + |
| 2 | -- | pExTra_67 | Amelie *67* replicate 2 | Non-toxic | + |
| 3 | -- | pExTra_67 | Amelie *67* replicate 3 | Non-toxic | + |

*Key: NG (no growth) - (no pink color) +(faint pink color) ++(obvious pink color) +++ (dark pink color)

Images taken after 5 days at 37 ºC

#### Gene *68*; Score *0*

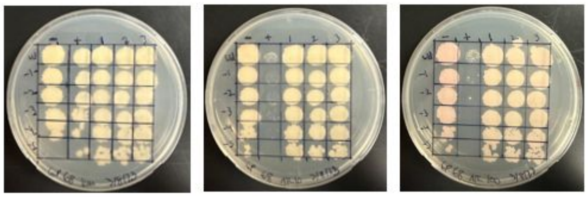


0 ng/ml aTc | 10 ng/ml aTc | 100 ng/ml aTc

| Lane | Gene ID | Plasmid name | Gene name | Toxic/Non-toxic | Colony color on 100 ng/ml aTc plate* |
|---|---|---|---|---|---|
| - Non-toxic control | -- | pExTra03 | Fruitloop *52* mutant | Non-toxic | + |
| + Toxic control | -- | pExTra02 | Fruitloop *52* | Toxic | - |
| 1 | | pExtra_68 | Amelie 68 replicate 1 | Non-toxic | + |
| 2 | | pExtra_68 | Amelie 68 replicate 2 | Non-toxic | + |
| 3 | | pExtra_68 | Amelie 68 replicate 3 | Non-toxic | + |

*Key: NG (no growth) - (no pink color) +(faint pink color) ++(obvious pink color) +++ (dark pink color)

Images taken after 5 days at 37 ºC

#### Gene *69*; Score *0*

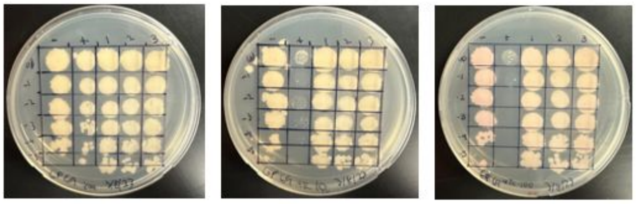


0 ng/ml aTc | 10 ng/ml aTc | 100 ng/ml aTc

| Lane | Gene ID | Plasmid name | Gene name | Toxic/Non-toxic | Colony color on 100 ng/ml aTc plate* |
|---|---|---|---|---|---|
| - Non-toxic control | -- | pExTra03 | Fruitloop *52* mutant | Non-toxic | + |
| + Toxic control | -- | pExTra02 | Fruitloop *52* | Toxic | - |
| 1 | | pExtra_69 | Amelie 69 replicate 1 | Non-toxic | + |
| 2 | | pExtra_69 | Amelie 69 replicate 2 | Non-toxic | + |
| 3 | | pExtra_69 | Amelie 69 replicate 3 | Non-toxic | + |

*Key: NG (no growth) - (no pink color) +(faint pink color) ++(obvious pink color) +++ (dark pink color)

Images taken after 7 days at 37 ºC

#### Gene *70*; Score *0*

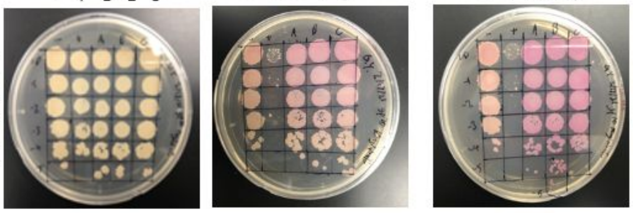


0 ng/ml aTc | 10 ng/ml aTc | 100 ng/ml aTc

| Lane | Gene ID | Plasmid name | Gene name | Toxic/Non-toxic | Colony color on 100 ng/ml aTc plate* |
|---|---|---|---|---|---|
| - Non-toxic control | -- | pExTra03 | Fruitloop *52* mutant | Non-toxic | ++ |
| + Toxic control | -- | pExTra02 | Fruitloop *52* | Toxic | - |
| 1 | -- | pExtra_70 | Amelie 70 replicate 1 | Non-toxic | +++ |
| 2 | -- | pExtra_70 | Amelie 70 replicate 2 | Non-toxic | +++ |
| 3 | -- | pExtra_70 | Amelie 70 replicate 3 | Non-toxic | +++ |

*Key: NG (no growth) - (no pink color) +(faint pink color) ++(obvious pink color) +++ (dark pink color)

Images taken after 5 days at 37 ºC

#### Gene *71*; Score *0*

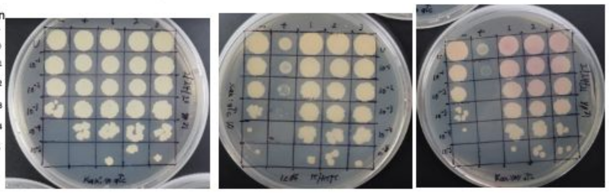


0 ng/ml aTc | 10 ng/ml aTc | 100 ng/ml aTc

| Lane | Gene ID | Plasmid name | Gene name | Toxic/Non-toxic | Colony color on 100 ng/ml aTc plate* |
|---|---|---|---|---|---|
| - Non-toxic control | -- | pExTra03 | Fruitloop *52* mutant | Non-toxic | + |
| + Toxic control | -- | pExTra02 | Fruitloop *52* | Toxic | - |
| 1 | -- | pExTra_71 | Amelie *71* replicate 1 | Non-toxic | + |
| 2 | -- | pExTra_71 | Amelie *71* replicate 2 | Non-toxic | + |
| 3 | -- | pExTra_71 | Amelie *71* replicate 3 | Non-toxic | + |

*Key: NG (no growth) - (no pink color) +(faint pink color) ++(obvious pink color) +++ (dark pink color)

Images taken after 5 days at 37 ºC

#### Gene *72*; Score *0*

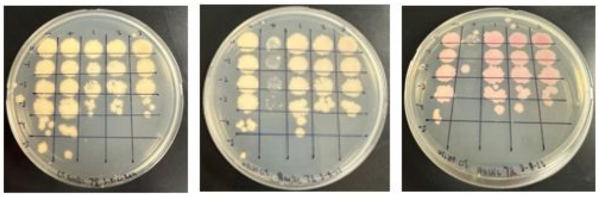


0 ng/ml aTc | 10 ng/ml aTc | 100 ng/ml aTc

| Lane | Gene ID | Plasmid name | Gene name | Toxic/Non-toxic | Colony color on 100 ng/ml aTc plate* |
|---|---|---|---|---|---|
| - Non-toxic control | -- | pExTra03 | Fruitloop *52* mutant | Non-toxic | + |
| + Toxic control | -- | pExTra02 | Fruitloop *52* | Toxic | - |
| 1 | | pExtra_72 | Amelie 72 replicate 1 | Non-toxic | ++ |
| 2 | | pExtra_72 | Amelie 72 replicate 2 | Non-toxic | ++ |
| 3 | | pExtra_72 | Amelie 72 replicate 3 | Non-toxic | ++ |

*Key: NG (no growth) - (no pink color) +(faint pink color) ++(obvious pink color) +++ (dark pink color)

Images taken after 5 days at 37 ºC

#### Gene *73*; Score *3*

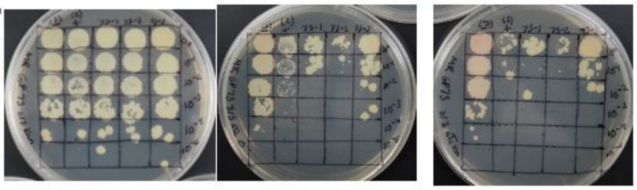


0 ng/ml aTc | 10 ng/ml aTc | 100 ng/ml aTc

| Lane | Gene ID | Plasmid name | Gene name | Toxic/Non-toxic | Colony color on 100 ng/ml aTc plate* |
|---|---|---|---|---|---|
| - Non-toxic control | -- | pExTra03 | Fruitloop *52* mutant | Non-toxic | + |
| + Toxic control | -- | pExTra02 | Fruitloop *52* | Toxic | - |
| 1 | | pExtra_73 | Amelie 73 replicate 1 | Toxic | - |
| 2 | | pExtra_73 | Amelie 73 replicate 2 | Toxic | - |
| 3 | | pExtra_73 | Amelie 73 replicate 3 | Toxic | - |

*Key: NG (no growth) - (no pink color) +(faint pink color) ++(obvious pink color) +++ (dark pink color)

#### Gene *74*; Score *1*

Images taken after <u>5</u> days at 37 ºC

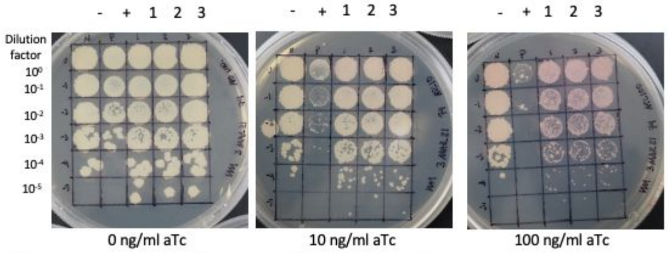


| Lane | Gene ID | Plasmid name | Gene name | Toxic/Non-toxic | Colony color on 100 ng/ml aTc plate* |
|---|---|---|---|---|---|
| - Non-toxic control | -- | pExTra03 | Fruitloop *52* mutant | Non-toxic | ++ |
| + Toxic control | -- | pExTra02 | Fruitloop *52* | Toxic | + |
| 1 | -- | pExTra_74 | Amelie *74*replicate 1 | Toxic | + |
| 2 | -- | pExTra_74 | Amelie *74* replicate 2 | Toxic | + |
| 3 | -- | pExTra_74 | Amelie *74* replicate 3 | Toxic | + |

*Key: NG (no growth) - (no pink color) +(faint pink color) ++(obvious pink color) +++ (dark pink color)

#### Gene *76*; Score *0*

Images taken after <u>5</u> days at 37 ºC

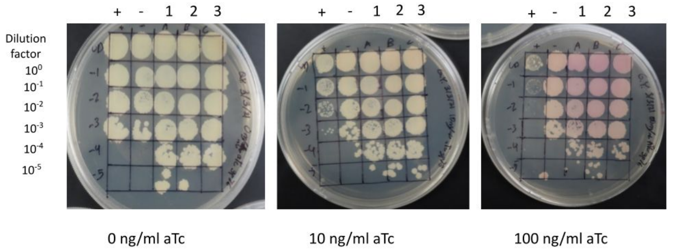


| Lane | Gene ID | Plasmid name | Gene name | Toxic/Non-toxic | Colony color on 100 ng/ml aTc plate* |
|---|---|---|---|---|---|
| - Non-toxic control | -- | pExTra03 | Fruitloop *52* mutant | Non-toxic | + |
| + Toxic control | -- | pExTra02 | Fruitloop *52* | Toxic | - |
| 1 | -- | pExTra_76 | Amelie 76 replicate 1 | Non-toxic | ++ |
| 2 | -- | pExTra_76 | Amelie 76 replicate 2 | Non-toxic | ++ |
| 3 | -- | pExTra_76 | Amelie 76 replicate 3 | Non-toxic | ++ |

*Key: NG (no growth) - (no pink color) +(faint pink color) ++(obvious pink color) +++ (dark pink color)

#### Gene *77*; Score *0*

Images taken after <u>5</u> days at 37 ºC

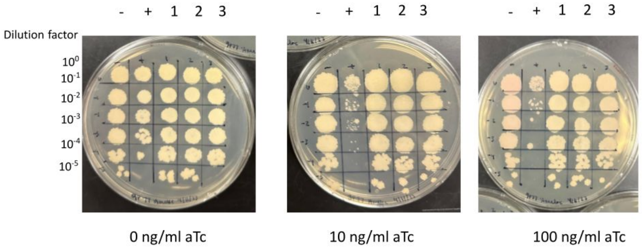


| Lane | Gene ID | Plasmid name | Gene name | Toxic/Non-toxic | Colony color on 100 ng/ml aTc plate* |
|---|---|---|---|---|---|
| - Non-toxic control | -- | pExTra03 | Fruitloop *52* mutant | Non-toxic | + |
| + Toxic control | -- | pExTra02 | Fruitloop *52* | Toxic | - |
| 1 | -- | pExTra_77 | Amelie *77* replicate 1 | Non-toxic | + |
| 2 | -- | pExTra_77 | Amelie *77* replicate 2 | Non-toxic | + |
| 3 | -- | pExTra_77 | Amelie *77* replicate 3 | Non-toxic | + |

*Key: NG (no growth) - (no pink color) +(faint pink color) ++(obvious pink color) +++ (dark pink color)

**Supplemental Figure 1**: Shown are the results of representative cytotoxicity assays for the 76 Amelie genes screened in this study. Each strain was spotted in triplicate alongside *M. smegmatis*/pExtra-Fruitloop 52 (+) and pExtra-Fruitloop52I170S (-) control strain in 7H10 Kan supplemented with 0, 10, or 100 ng/ml aTc. In all experiments, $10^0$ to $10^{-5}$ dilutions are shown. Plates were monitored over 4 or 5 days at 37℃, with results shown to best illustrate effects on colony color and size. Colony color was scored using the indicated key shown at the bottom of the data card.

**Supplemental Table 1: Primer sequences used in this study to amplify genes**

| Primer Name | Primer Sequence (5’ to 3’) |
|---|---|
| oAmelie1 _F | atgcggaggaatcacttccatATGTCGCTCGCGCCC |
| oAmelie1 _R | tgcaggatccgactcgagtgtcgacTCAGCCGGAAGGTGCGC |
| oAmelie2 _F | atgcggaggaatcacttccatATGCGCGCACCCGCTAC |
| oAmelie2 _R | tgcaggatccgactcgagtgtcgacTCACTCATCGCGGCCAC |
| oAmelie3 _F | atgcggaggaatcacttccatATGGCCGCGATGAGTGAC |
| oAmelie3 _R | tgcaggatccgactcgagtgtcgacTCAGAGGTCCGAGCCATC |
| oAmelie4 _F | atgcggaggaatcacttccatATGGCTCGGACCTCTGAG |
| oAmelie4 _R | tgcaggatccgactcgagtgtcgacTCACACGAACATTGCGCC |
| oAmelie5 _F | atgcggaggaatcacttccatATGATCCCGCAGCCTTTCG |
| oAmelie5 _R | tgcaggatccgactcgagtgtcgacTCACTGAGCCGAACCAGC |
| oAmelie6 _F | atgcggaggaatcacttccatATGACGGAACCCAAGGC |
| oAmelie6 _R | tgcaggatccgactcgagtgtcgacTCACTGCAGGAAGTCCTC |
| oAmelie7 _F | atgcggaggaatcacttccatATGAACCGCGACGAGCTG |
| oAmelie7 _R | tgcaggatccgactcgagtgtcgacTCACTCAGCCTGTGCCTG |
| oAmelie8 _F | atgcggaggaatcacttccatATGGCTGATAACGCGAGCG |
| oAmelie8 _R | tgcaggatccgactcgagtgtcgacTCAGTCGCCGGAACGCAG |
| oAmelie9 _F | atgcggaggaatcacttccatATGGCTGACATTTCACGCTCC |
| oAmelie9 _R | tgcaggatccgactcgagtgtcgacTCAGTCCCCTGCCGGGAG |
| oAmelie10 _F | atgcggaggaatcacttccatATGCTGGCGACGCTGGAC |
| oAmelie10 _R | tgcaggatccgactcgagtgtcgacTCAGTACCTCTCGCTGCCC |
| oAmelie11 _F | atgcggaggaatcacttccatATGTTCCCGACGCCCCAC |
| oAmelie11 _R | tgcaggatccgactcgagtgtcgacTCAAGCGGTCGGTATGGC |
| oAmelie12 _F | atgcggaggaatcacttccatATGCCATACCGACCGCTTG |
| oAmelie12 _R | tgcaggatccgactcgagtgtcgacTCACCGTGGCCCCAACTC |
| oAmelie13 _F | atgcggaggaatcacttccatATGAGCGTGCTCGTTCCGC |
| oAmelie13 _R | tgcaggatccgactcgagtgtcgacTCAGCTGCGGCGCTCGG |
| oAmelie14 _F | atgcggaggaatcacttccatATGGCAGACGAGCCTACTTC |
| oAmelie14 _R | tgcaggatccgactcgagtgtcgacTCAGACGCCGATCTTCTGAC |
| oAmelie15 _F | atgcggaggaatcacttccatATGACTGAGACCAACGTCGACAGC |
| oAmelie15 _R | tgcaggatccgactcgagtgtcgacTCACCTGCCGCGCCGCC |

| | |
|---|---|
| oAmelie16 _F | atgcggaggaatcacttccatATGACTGAGACCAACGTCGAC |
| oAmelie16 _R | tgcaggatccgactcgagtgtcgacTCACGCCCCCTTGCTCTTG |
| oAmelie17 _F | atgcggaggaatcacttccatATGAGCGCTACGTACTACCTC |
| oAmelie17 _R | tgcaggatccgactcgagtgtcgacTCACCTGTATCGAGCGCGAG |
| oAmelie18 _F | atgcggaggaatcacttccatATGGACAAGAAGTACACGGGC |
| oAmelie18 _R | tgcaggatccgactcgagtgtcgacTCACGACCACGCCATCCG |
| oAmelie19 _F | atgcggaggaatcacttccatATGTACGTGAAAGATGGCCGC |
| oAmelie19 _R | tgcaggatccgactcgagtgtcgacTCAGAACATGTCTCCTGATCCG |
| oAmelie20 _F | atgcggaggaatcacttccatATGGAAATGCCCACACTGCCAC |
| oAmelie20 _R | tgcaggatccgactcgagtgtcgacTCACTCGTCGAAAGTGATGGAC |
| oAmelie21 _F | atgcggaggaatcacttccatATGGCCGAGCTTGCGC |
| oAmelie21 _R | tgcaggatccgactcgagtgtcgacTCACTCCCCTTGCGGCAC |
| oAmelie22 _F | atgcggaggaatcacttccatATGCCGTACACCAAGAGTTACC |
| oAmelie22 _R | tgcaggatccgactcgagtgtcgacTCACTTACCTCCTGCCTTGC |
| oAmelie23 _F | atgcggaggaatcacttccatATGCCTCCCGTCTACGACC |
| oAmelie23 _R | tgcaggatccgactcgagtgtcgacTCACGTCGTCGCCGTGAC |
| oAmelie24 _F | atgcggaggaatcacttccatATGGCGTGGTCAACAAACCC |
| oAmelie24 _R | tgcaggatccgactcgagtgtcgacTCACTGGTAGAACCTCAGCCAG |
| oAmelie25 _F | atgcggaggaatcacttccatATGGCAGGAGCGGCTG |
| oAmelie25 _R | tgcaggatccgactcgagtgtcgacTCAGTCCGTCAGGTCGTAGATG |
| oAmelie26 _F | atgcggaggaatcacttccatATGATTATCTACCGCGCTAACGTC |
| oAmelie26 _R | tgcaggatccgactcgagtgtcgacTCACGCGCTCACCCC |
| oAmelie27 _F | atgcggaggaatcacttccatATGAGCAAGCCTGTTCTGCTC |
| oAmelie27 _R | tgcaggatccgactcgagtgtcgacTCACGCCGCCATCGC |
| oAmelie28 _F | atgcggaggaatcacttccatATGGCGAAATCGGCCGC |
| oAmelie28 _R | tgcaggatccgactcgagtgtcgacTCACCGCTCGACCGCC |
| oAmelie29 _F | atgcggaggaatcacttccatATGAGCGCCGACACGTTG |
| oAmelie29 _R | tgcaggatccgactcgagtgtcgacTCATTCGATCGCCCCGTCG |
| oAmelie30 _F | atgcggaggaatcacttccatATGAGCCTCGCAGAACG |
| oAmelie30 _R | tgcaggatccgactcgagtgtcgacTCAGACAGCGACACGGG |
| oAmelie31 _F | atgcggaggaatcacttccatATGTCGCTGTCTGACCGAC |
| oAmelie31 _R | tgcaggatccgactcgagtgtcgacTCACACGACGCTGAGGC |

| | |
|---|---|
| oAmelie32 _F | atgcggaggaatcacttccatATGAACGCTCACACAATGACAG |
| oAmelie32 _R | tgcaggatccgactcgagtgtcgacTCACCGATCCTTCTTGCCC |
| oAmelie33 _F | atgcggaggaatcacttccatATGGCCGGGAGTGATGTTC |
| oAmelie33 _R | tgcaggatccgactcgagtgtcgacTCAGTGATCGCAGACGTCG |
| oAmelie34 _F | atgcggaggaatcacttccatATGAGCGAGTACACCAAGGAC |
| oAmelie34 _R | tgcaggatccgactcgagtgtcgacTCACTTGTTCTTTGGCAGCTTG |
| oAmelie35 _F | atgcggaggaatcacttccatATGATCGAGCACTTTTACCTCGG |
| oAmelie35 _R | tgcaggatccgactcgagtgtcgacTCAGGCGACGTCGAACAAC |
| oAmelie36 _F | atgcggaggaatcacttccatATGGCATCGCTTCGCACC |
| oAmelie36 _R | tgcaggatccgactcgagtgtcgacTCAGTTCCGGTCGAGTTGG |
| oAmelie37 _F | atgcggaggaatcacttccatATGCTCGACGATCTCAATAGAATC |
| oAmelie37 _R | tgcaggatccgactcgagtgtcgacTCAGTGGGTTGACTGCAGG |
| oAmelie38 _F | atgcggaggaatcacttccatATGCCAGACCATGACGACG |
| oAmelie38 _R | tgcaggatccgactcgagtgtcgacTCACCGCAGAATGGCGTCC |
| oAmelie39 _F | atgcggaggaatcacttccatATGCAGAACTTTAGACACGAACTGC |
| oAmelie39 _R | tgcaggatccgactcgagtgtcgacTCATGCGGCGGCCCG |
| oAmelie40 _F | atgcggaggaatcacttccatATGAGTTCTCCCGCAACCG |
| oAmelie40 _R | tgcaggatccgactcgagtgtcgacTCAACTGCATGAGCGGCTC |
| oAmelie41 _F | atgcggaggaatcacttccatATGCAGTTAGCCCGCCAC |
| oAmelie41 _R | tgcaggatccgactcgagtgtcgacTCATTTCTCGGCTCCCCC |
| oAmelie42 _F | atgcggaggaatcacttccatATGCCGAGCAGAATCTTAGCG |
| oAmelie42 _R | tgcaggatccgactcgagtgtcgacTCATCGAGCGACCGCC |
| oAmelie43 _F | atgcggaggaatcacttccatATGACCCCGGCGCCG |
| oAmelie43 _R | tgcaggatccgactcgagtgtcgacTCACCAAGGCAGCACCC |
| oAmelie44 _F | atgcggaggaatcacttccatATGGTGACCCTGACTCACG |
| oAmelie44 _R | tgcaggatccgactcgagtgtcgacTCACAGCATCACCGTCCG |
| oAmelie45 _F | atgcggaggaatcacttccatATGACGGTTCGGCGCATC |
| oAmelie45 _R | tgcaggatccgactcgagtgtcgacTCACGCTGCGCCCTCG |
| oAmelie46 _F | atgcggaggaatcacttccatATGACCGGCGGCCCATTC |
| oAmelie46 _R | tgcaggatccgactcgagtgtcgacTCACGCGACCACCTCC |
| oAmelie47 _F | atgcggaggaatcacttccatATGACCGACCGGCCGC |
| oAmelie47 _R | tgcaggatccgactcgagtgtcgacTCATTCAGTTGTGCTCAGCGC |

| | |
|---|---|
| oAmelie48 _F | atgcggaggaatcacttccatATGGCTGACGTTTCGGACC |
| oAmelie48 _R | tgcaggatccgactcgagtgtcgacTCATCCCTCTTCTTCGACCAG |
| oAmelie49 _F | atgcggaggaatcacttccatATGAGCGCAAGGAATCTGATCG |
| oAmelie49 _R | tgcaggatccgactcgagtgtcgacTCACAGGTGCGCCCC |
| oAmelie50 _F | atgcggaggaatcacttccatATGAGCGCCAATGCGAAGTTC |
| oAmelie50 _R | tgcaggatccgactcgagtgtcgacTCACGCGACGTCTCCC |
| oAmelie51 _F | atgcggaggaatcacttccatATGAGGCGCCCGGCGC |
| oAmelie51 _R | tgcaggatccgactcgagtgtcgacTCAGGCCGCGGATTCGC |
| oAmelie52 _F | atgcggaggaatcacttccatATGCGCAAGGTAATTGCCGTG |
| oAmelie52 _R | tgcaggatccgactcgagtgtcgacTCACAGCAAGATGAAGCTGCC |
| oAmelie53 _F | atgcggaggaatcacttccatATGAGCCGCCATAACTGCAG |
| oAmelie53 _R | tgcaggatccgactcgagtgtcgacTCACGTCCCGTCGTAGAAAC |
| oAmelie54 _F | atgcggaggaatcacttccatATGAGCAATGACTCGTACGACTTC |
| oAmelie54 _R | tgcaggatccgactcgagtgtcgacTCACTGCTTCGCTGCCAG |
| oAmelie55 _F | atgcggaggaatcacttccatATGATCACGATTTACACGACCG |
| oAmelie55 _R | tgcaggatccgactcgagtgtcgacTCACGCTGCCTGCCG |
| oAmelie56 _F | atgcggaggaatcacttccatATGAACACGACGGCCCTG |
| oAmelie56 _R | tgcaggatccgactcgagtgtcgacTCAAAGGGTTGGGCGGC |
| oAmelie57 _F | atgcggaggaatcacttccatATGAACGGACTTTCTGACCTGC |
| oAmelie57 _R | tgcaggatccgactcgagtgtcgacTCACCGCCGGAAACCG |
| oAmelie58 _F | atgcggaggaatcacttccatATGCACCTCGACCACACAAC |
| oAmelie58 _R | tgcaggatccgactcgagtgtcgacTCAGGCATCCCCAGCC |
| oAmelie59 _F | atgcggaggaatcacttccatATGCCTGATCTGATCGAGTTG |
| oAmelie59 _R | tgcaggatccgactcgagtgtcgacTCACTTCGGCGACACCTC |
| oAmelie60 _F | atgcggaggaatcacttccatATGACTTTGCCGGTTTCTGTGG |
| oAmelie60 _R | tgcaggatccgactcgagtgtcgacTCATCCCATCACCTGACTAGG |
| oAmelie61 _F | atgcggaggaatcacttccatATGTACACAGAGGCATGGTTGTCG |
| oAmelie61 _R | tgcaggatccgactcgagtgtcgacTCACGCCAGCGCCGC |
| oAmelie62 _F | atgcggaggaatcacttccatATGAAGCGTATGAAGGCGTTTCG |
| oAmelie62 _R | tgcaggatccgactcgagtgtcgacTCAGGCGACGTTCGCCC |
| oAmelie63 _F | atgcggaggaatcacttccatATGAAGGTCAGCAAGGTTCG |
| oAmelie63 _R | tgcaggatccgactcgagtgtcgacTCAGGCGCCGATGCTG |

| | |
|---|---|
| oAmelie64 _F | atgcggaggaatcacttccatATGAACACTCTTCACCTCACGG |
| oAmelie64 _R | tgcaggatccgactcgagtgtcgacTCATCGGGCGACCTCGC |
| oAmelie65 _F | atgcggaggaatcacttccatATGAGCGCCGAGGCC |
| oAmelie65 _R | tgcaggatccgactcgagtgtcgacTCACGCAACCACCTCCG |
| oAmelie66 _F | atgcggaggaatcacttccatATGAGCGCCGCCGAGG |
| oAmelie66 _R | tgcaggatccgactcgagtgtcgacTCACACCCAGCCGGGCAC |
| oAmelie67 _F | atgcggaggaatcacttccatATGTCGCCGACGGTAATCAAC |
| oAmelie67 _R | tgcaggatccgactcgagtgtcgacTCACAGCCCCTTCACGTTG |
| oAmelie68 _F | atgcggaggaatcacttccatATGGCCGACTATTCGTTCGC |
| oAmelie68 _R | tgcaggatccgactcgagtgtcgacTCACCCATGCCGCAGCAC |
| oAmelie69 _F | atgcggaggaatcacttccatATGGGTAGTGCGCCGGTG |
| oAmelie69 _R | tgcaggatccgactcgagtgtcgacTCACTCATCAGCCACCACC |
| oAmelie70 _F | atgcggaggaatcacttccatATGAGTGATGTTGTTGATACGCG |
| oAmelie70 _R | tgcaggatccgactcgagtgtcgacTCAGCCAAACAGGCGCC |
| oAmelie71 _F | atgcggaggaatcacttccatATGTTCAAGATGATCGTTCAGCTTC |
| oAmelie71 _R | tgcaggatccgactcgagtgtcgacTCACGCGGCCGCCAG |
| oAmelie72 _F | atgcggaggaatcacttccatATGGACGCTCCAACGCATTTC |
| oAmelie72 _R | tgcaggatccgactcgagtgtcgacTCAGTCGGTGTACTTCACCTCG |
| oAmelie73 _F | atgcggaggaatcacttccatATGCCGAAACGTACCGAGG |
| oAmelie73 _R | tgcaggatccgactcgagtgtcgacTCAGCGCCACCAGCC |
| oAmelie74 _F | atgcggaggaatcacttccatATGATGGTTAAGGGTATCGAGTGG |
| oAmelie74 _R | tgcaggatccgactcgagtgtcgacTCAGCAGCCGTGCACCAG |
| oAmelie75 _F | atgcggaggaatcacttccatATGTCGACCACCTTCAAGTACC |
| oAmelie75 _R | tgcaggatccgactcgagtgtcgacTCAGTCCTCGGCCGC |
| oAmelie76 _F | atgcggaggaatcacttccatATGTCCGAGAACGCTCCC |
| oAmelie76 _R | tgcaggatccgactcgagtgtcgacTCACCAGTCGTACTGCTGGTAG |
| oAmelie77 _F | atgcggaggaatcacttccatATGACGGCGGGCCGC |
| oAmelie77 _R | tgcaggatccgactcgagtgtcgacTCAGGGTCGCCACGTCC |
| pExTra_F | GTCGACACTCGAGTCGGATCCTG |
| pExTra_R | ATGGAAGTGATTCCTCCGCATGC |
| pExTra_seqF | GTACCCGTGTGTACGACCAGC |
| pExTra_uniR | CCCTTCGAGACCATAGATCTGTTCC |

| | |
|---|---|
| oAmelie_6ia_seq | ggacacgatcactgcaagtgc |
| oAmelie_6ib_seq | gtttcgcacgcttgggtg |
| oAmelie_17ia_seq | AGCGTTCGGGCTCGACG |
| oAmelie_17ib_seq | AGCGAGTCATTGACTTCATGGGG |
| oAmelie_17ic_seq | GTTCCTCATCGGCCTGCATC |
| oAmelie_17id_seq | TGGCGAACCGAGAACCGAAC |
| oAmelie_17ie_seq | CAGTGGTTGTGGAATGCGGC |
| oAmelie_23ia_seq | CCTGACCGCAGCCAACG |
| oAmelie_23ib_seq | CCATCCCGAAGCCCCAG |
| oAmelie_23ic_seq | GGACGTGCACCAGATCCTC |
| oAmelie_24ia_seq | GTCTGCTGCTCAACGGTCG |
| oAmelie_57ia_seq | CCGGGCAGCTACAACATGAAG |
| oAmelie_57ib_seq | CATGTACCGGCCGCTGAG |
